# Evolutionary repair: changes in multiple functional modules allow meiotic cohesin to support mitosis

**DOI:** 10.1101/844423

**Authors:** Yu-Ying Phoebe Hsieh, Vasso Makrantoni, Daniel Robertson, Adèle L Marston, Andrew W Murray

**Affiliations:** Department of Molecular and Cellular Biology, Harvard University, Cambridge MA, 02138; The Wellcome Centre for Cell Biology, University of Edinburgh, Edinburgh EH9 3BF, UK

## Abstract

Different members of the same protein family often perform distinct cellular functions. How much are these differing functions due to changes in a protein’s biochemical activity versus changes in other proteins? We asked how the budding yeast, *Saccharomyces cerevisiae,* evolves when forced to use the meiosis-specific kleisin, Rec8, instead of the mitotic kleisin, Scc1, during the mitotic cell cycle. This perturbation impairs sister chromosome linkage and reduces reproductive fitness by 45%. We evolved 15 populations for 1750 generations, substantially increasing their fitness, and analyzed their genotypes and phenotypes. We found no mutations in Rec8, but many populations had mutations in the transcriptional mediator complex, cohesin-related genes, and cell cycle regulators that induce S phase. These mutations improve sister chromosome cohesion and slow genome replication in Rec8-expressing cells. We conclude that changes in known and novel partners allow proteins to improve their ability to perform new functions.

## Introduction

How does natural selection change a protein’s function during evolution? The biological function of a protein is determined by its intrinsic biochemical activity and its interactions with other proteins that control its abundance, activity, and location within the cell. Paralogs, proteins which arise by gene duplication and perform different functions are good candidates for studying how evolution modifies protein function (Orengo and Thornton, 2005; Chothia et al., 2003). Paralogs can diverge in many ways. Changes in their promoters and enhancers lead to altered patterns of gene expression (Gagnon-Arsenault et al., 2013; Hittinger and Carroll, 2007) and changes in their amino acid sequences alter biochemical activity (Voordeckers et al., 2012), patterns of post-translational modification (Amoutzias et al., 2010; Nguyen Ba et al., 2014), or the identity of interacting partners (Aakre et al., 2015). It is much less easy to identify the changes in other genes that collaborate to increase the functional divergence between paralogs. One approach to this problem is to ask what must change, either in the candidate protein or elsewhere in the genome, to allow a protein to perform the function of a paralog from which it diverged hundreds of millions of years ago.

To study how protein function evolves, we studied the kleisin protein family whose members organize the structure of chromosomal DNA. In prokaryotes and eukaryotes, kleisins bind to SMC (structural maintenance complex) proteins to form a ring complex (Schleiffer et al., 2003) that interacts with chromosomes. In most bacteria and archaea, there is a single kleisin and SMC protein (Melby et al., 1998; Soppa, 2001). In the eukaryotes, kleisin and SMC proteins have duplicated and acquired specialized functions (Cobbe and Heck, 2004; Schleiffer et al., 2003). Kleisin-γ proteins associate with Smc2/Smc4 heterodimers to form the condensin complex (Hirano, 2012), which regulates chromosome structure in mitosis and meiosis. Kleisin-α proteins interact with Smc1/Smc3 heterodimers to form cohesin, the complex that holds sister chromosomes together (Nasmyth, 2002) and regulates the timing of chromosome segregation: cohesin holds chromosomes together from S phase, when DNA replication occurs, until the proteolytic cleavage of kleisn by separase opens the ring and allows sister chromosomes to separate from each other at anaphase.

Most eukaryotes have two different kleisin-α proteins. The mitotic kleisin holds sister chromosomes together in mitosis, whereas the meiotic kleisin is expressed only in meiosis (Mehta et al., 2012). Both kleisins interact with Smc1 and Smc3 but their proteolysis is regulated differently to produce the different patterns of chromosome segregation in mitosis and meiosis. In mitosis, the protease that cleaves kleisin is activated at anaphase leading to sister chromosome separation (Marston, 2014; Uhlmann et al., 1999; Uhlmann et al., 2000). In meiosis, however, the regulation of kleisin cleavage is modified to allow two rounds of chromosome segregation to follow a single round of DNA replication, producing four haploid genomes: in meiosis I, cleaving the kleisin on the chromosome arms allows homologous chromosomes to segregate from each other, and then in meiosis II, cleaving the remaining kleisin, near the centromeres, allows sister centromeres to segregate from each other (Buonomo et al., 2000; Marston, 2014). Both cohesin complexes have additional functions. Mitotic cohesin regulates chromosome condensation (Guacci et al., 1997; Lazar-Stefanita et al., 2017), gene expression (Donze et al., 1999), and DNA damage repair (Heidinger-Pauli et al., 2008; Wu and Yu, 2012) and meiotic cohesin regulates meiotic recombination (Brar et al., 2009; Klein et al., 1999) and chromosome topology (Schalbetter et al., 2019). Most eukaryotes have both mitotic and meiotic kleisins, suggesting that the duplication and divergence of these paralogous proteins occurred at or soon after the evolutionary origin of the eukaryotes (Dorsett and Merkenschlager, 2013; Feeney et al., 2010; Peric-Hupkes and van Steensel, 2008).

In the budding yeast, *Saccharomyces cerevisiae*, the functions of the mitotic and meiotic kleisins are not interchangeable. In this species, the mitotic kleisin is encoded by *SCC1*; the meiotic kleisin is encoded by *REC8*. Replacing the coding sequence of *REC8* by that of *SCC1* during meiosis disrupts meiotic chromosome segregation (Brar et al., 2009; Toth et al., 2000) and the opposite experiment, expressing *REC8* from the *SCC1* promoter in mitosis, slows cell proliferation (Buonomo et al., 2000). These results suggest that the functional difference between yeast kleisin proteins is mainly determined by the difference between their amino acid sequences rather than the promoters that control their expression. The ancient evolutionary separation of the mitotic and meiotic kleisins makes it hard to answer two questions: 1) which mutations in kleisin produced changes in its function rather than accumulating because of other selective forces or genetic drift and 2) what fraction of the mutations that altered kleisin function occurred in other proteins rather than kleisin itself.

We used experimental evolution to ask how cells adapt when they are selected to use one protein for a function that is normally performed by its paralog. We substituted the budding yeast meiotic kleisin, Rec8, for its mitotic counterpart, Scc1, thus requiring Rec8 to support mitotic rather than meiotic chromosome segregation. Can cells evolve to proliferate faster and more accurately while using Rec8 for mitosis? If so, do the adaptive mutations occur in kleisin or elsewhere in the genome? We asked these questions by evolving parallel yeast populations that expressed Rec8 in place of Scc1 for 1750 mitotic generations. We recovered no mutations in Rec8, but found adaptive mutations in the transcriptional mediator complex, cell cycle regulators that induce the G1-to-S transition, and cohesin-related genes; these mutations restore sister chromosome cohesion and thus increase the fitness of the evolved populations. Unexpectedly, we found that replacing Scc1 with Rec8 leads to earlier firing of replication origins. All three classes of adaptive mutations restored the timing of genome replication to the wild-type pattern. Engineering mutations that reduced replication origin firing or slowed replication forks improved the fitness of Rec8-dependent cells, revealing a new link between genome replication and sister chromosome cohesion. Our results suggest that the fastest way of adapting a protein to a novel function can be to modify the partners it directly or indirectly interacts with, rather than modifying the protein that must acquire the new function

## Results

### Using the meiotic kleisin, Rec8, for mitosis leads to multiple defects

We examined the consequences of replacing Scc1 with Rec8 in the mitotic cell cycle (Fig. 1A). Previous studies showed that replacing Scc1 with Rec8 impairs mitotic growth (Buonomo et al., 2000) and DNA damage repair (Heidinger-Pauli et al., 2008), showing that Rec8 cannot completely substitute for Scc1 in mitosis. We compared the reproductive fitness and the cellular and molecular phenotypes of the Rec8- and Scc1-expressing strains, referring to the Scc1-expresssing strain as the wild type. We used competitive growth to measure the fitness of the Rec8-expressing strain relative to wild type in rich media: the fitness of the Rec8-expressing strain is only 55% that of wild type (Fig. 1B).

**Figure 1.**
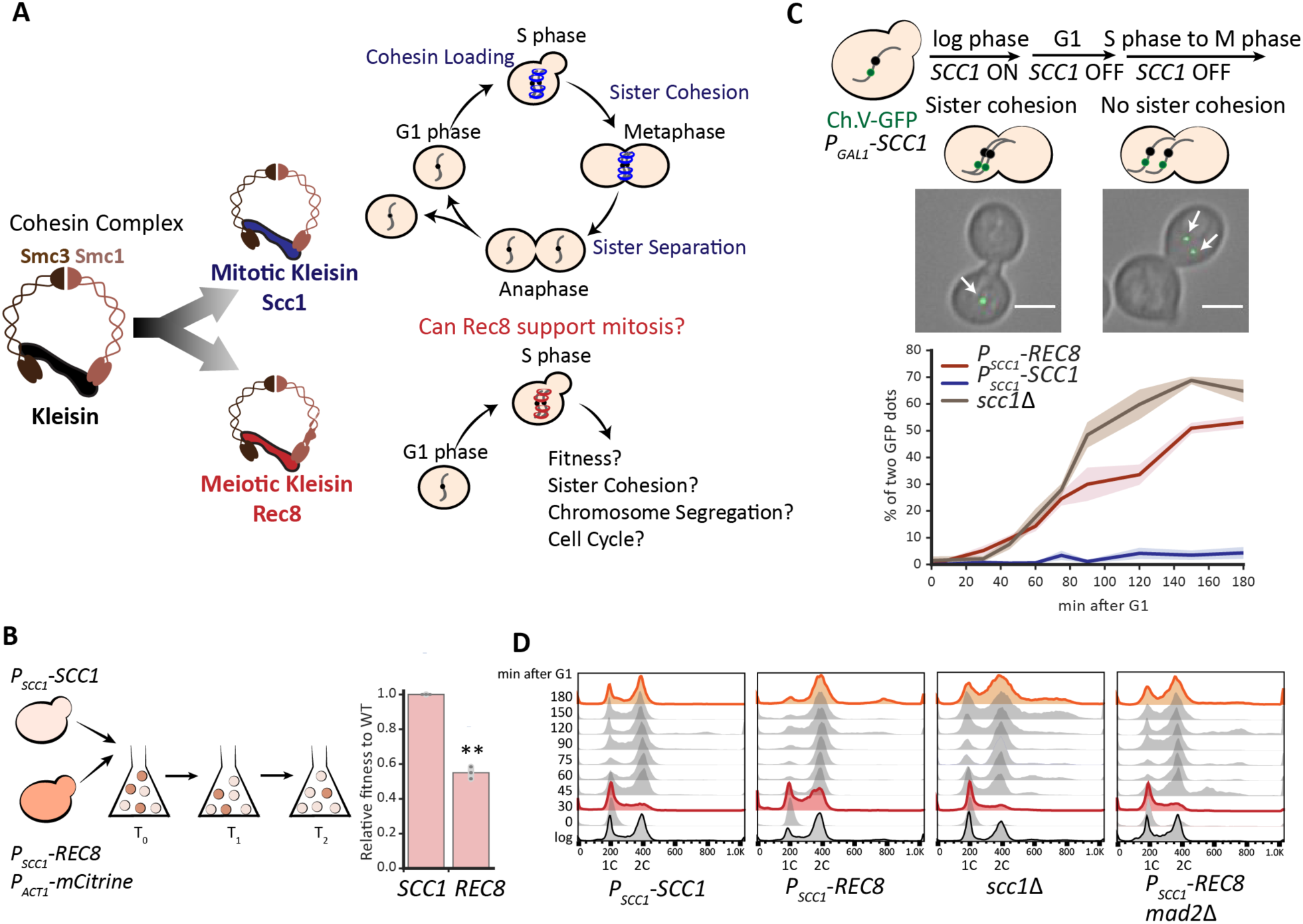
Expressing Rec8 in place of Scc1 impairs the mitotic cell cycle and sister chromosome cohesion. **(A)** A diagram of the mitotic and meiotic cohesin complexes. Mitotic cohesin holds replicated sister chromosomes together in mitosis. We investigated the ability of the meiotic cohesin to support mitosis. **(B)** The fitness of Rec8-expressing cells is 55% of a wild-type strain expressing Scc1. The Rec8-expressing strain expressed a fluorescent marker (*P_ACT1_*-*mCitrine*) and was competed against the wild type. The fitness of the Rec8-expressing cells relative to wild type was calculated as changes in the ratio of these two strains over multiple generations. In the right panel, the darker gray points represent the values of three biological replicates and the thinner gray bar represents one standard deviation on each side of the mean of these measurements. (two-tailed Student *t* test, ** *p* < 0.01) **(C)** The Rec8-expressing strain cannot maintain sister chromosome cohesion in mitosis. All the strains (*P_SCC1_*-*REC8*, *P_SCC1_*-*SCC1*, and *scc1*Δ) carried a *P_GAL1_*-*SCC1* copy integrated in the genome to allow the acute effect of altered kleisin expression to be analyzed. To examine sister chromosome cohesion in a single cell cycle, Scc1 expression was switched off in G1-arrested populations by transferring cells to YEP containing 2% Raffinose and alpha-factor. Cells were released to YPD containing benomyl to resume cell cycle and held in mitosis. Two different patterns of sister chromosome cohesion are shown: A budded cell with a single GFP dot represents functional sister chromosome cohesion; a budded cell with two GFP dots represents lack of sister chromosome cohesion. A single GFP dot is marked with a white arrow. At least 100 cells were imaged at each time point in each experiment. Three biological repeats were performed at each time point and for each strain; the right panel showed mean and standard deviation for the wild type (blue), Rec8-expressing (red), and *scc1*Δ (brown) strains. The scale bar is 5μm. **(D)** The Rec8-expressing strain progressed through S phase faster and mitosis slower. All strains were cultured as in Fig. 1C but cells were released in YPD to allow completion of the first cell cycle and entry into the second. Samples were collected at the indicated timepoints to examine DNA content by flow cytometry. Cell cycle profiles at 30 and 180 minutes are labeled in red and orange respectively.

We examined sister chromosome cohesion and chromosome segregation in Rec8-expressing cells. Because cells mis-segregating chromosomes become progressively more aneuploid, we wanted to examine acute rather than chronic effects of replacing Scc1 with Rec8. We expressed *REC8* from the endogenous *SCC1* promoter and conditionally expressed an additional copy of *SCC1* from the *GAL1* promoter. The *GAL1* promoter is rapidly repressed by glucose (Flick and Johnston, 1990; Johnston et al., 1994), allowing us to repress *SCC1* expression rapidly and study the function of Rec8 in a single mitotic cell cycle. We confirmed that when *SCC1* is turned off, expressing Rec8 slows progress through the cell cycle (Fig. S1A). We assayed cohesion between sister chromosomes by following a single, GFP-tagged chromosome through mitosis. Chromosome V was labeled by the binding of a GFP-*tet* repressor fusion to an array of *tet* operators integrated near the centromere (Michaelis et al., 1997; Uhlmann and Nasmyth, 1998). We asked if Rec8 could hold sister chromosomes together from S phase to mitosis by following the GFP-labeled centromeres (henceforth GFP dots) under the microscope as cells were released from a G1 arrest, allowed to proceed synchronously through the cell cycle, and then arrested in mitosis by benomyl that depolymerizes microtubules. In this assay, a pair of linked sister chromosomes appears as a single GFP dot whereas sister chromosomes that have lost cohesion appear as two GFP dots (Fig. 1C). The fraction of cells with two GFP dots in a population indicates the degree to which sister chromosomes have separated. From S phase to mitosis, the majority of the wild-type population showed a single GFP dot as expected (Fig. 1C). In the Rec8-expressing strain, 10% of the population showed two GFP dots during S phase and this fraction rose to 50% in mitosis (Fig. 1C). During a single cell cycle, the defect in sister chromosome cohesion of the Rec8-expressing strain was smaller than the defect in a strain completely lacking Scc1 (Fig. 1C), showing that Rec8 retains some cohesin function.

Sister chromosomes must be linked to each other to allow their kinetochores to attach stably to opposite poles of the mitotic spindle. In this orientation, forces exerted on the kinetochores create tension that inactivates the spindle checkpoint, leading to activation of the anaphase-promoting complex, the activation of separase, the cleavage of kleisin, and the onset of anaphase (Marston, 2014). The Rec8-expressing strain frequently failed to correctly orient sister kinetochores (Fig. S2), a defect likely to cause errors in chromosome segregation (Fig. S1B). We hypothesized that the sister chromosome cohesion defect would activate the spindle checkpoint, thus prolonging mitosis. By tracking a G1-synchronized population, we found that the Rec8-expressing strain accumulated more cells with a 2C DNA content compared to wild type (Fig. 1D). Removing Mad2, a spindle checkpoint protein (Li and Murray, 1991; Shah and Cleveland, 2000), from the Rec8-expressing strain, increases the fraction of cells with a 1C DNA content (from 90 to 180 minutes, Fig. 1D), suggesting that the sister chromosome cohesion defect activates the spindle checkpoint. In addition, the Rec8-expressing strain showed a shorter S phase than the wild-type. This phenotype is not due to faster escape from a G1 arrest since budded cells accumulate with indistinguishable frequency in wild-type and Rec8-expressing strains (Fig. S3). Unexpectedly, the *mad2*Δ, Rec8-expressing strain progressed through S phase with similar kinetics to wild type (see Discussion). In cells lacking kleisin or the cohesin loading complex mutant, the timing of genome replication is identical to wild-type (Uhlmann and Nasmyth, 1998), suggesting that the accelerated genome replication is a specific feature of Rec8-expressing cells. In summary, replacing Scc1 with Rec8 leads to profound defects in sister cohesion and accelerates genome replication by an unknown mechanism.

We asked whether the phenotype of Rec8-expressing cells could be explained by reduced cohesin levels or reduced cohesin binding to mitotic chromosomes. We measured Rec8 levels in a synchronous cell cycle and compared them with those of Scc1 (Fig. 2A). Scc1 was barely detectable in G1, peaked during S phase, and its cleavage product was detected at 60 minutes as cells entered anaphase. During G1 and S phase, there was four-fold less Rec8 than Scc1. At 90 minutes, the Rec8 protein level decreased but we did not detect the cleavage product of Rec8, either because the onset of anaphase was asynchronous or the cleavage product is too unstable to be detected (Buonomo et al., 2000). The lower Rec8 protein level could be due to inefficient protein synthesis or protein instability. We tested the second hypothesis by examining the stability of Scc1 and Rec8 in mitotically-arrested cells: the half-life of Scc1 is 200 minutes whereas Rec8 has a half-life of only 58 minutes (Fig. 2B). The instability of Rec8 is due to the weak separase activity that exists outside anaphase (Uhlmann et al., 1999): reducing separase activity with a temperature sensitive mutation, *esp1-1*, (Ho et al., 2015) increases the half-life of Rec8 to 190 minutes.

**Figure 2.**
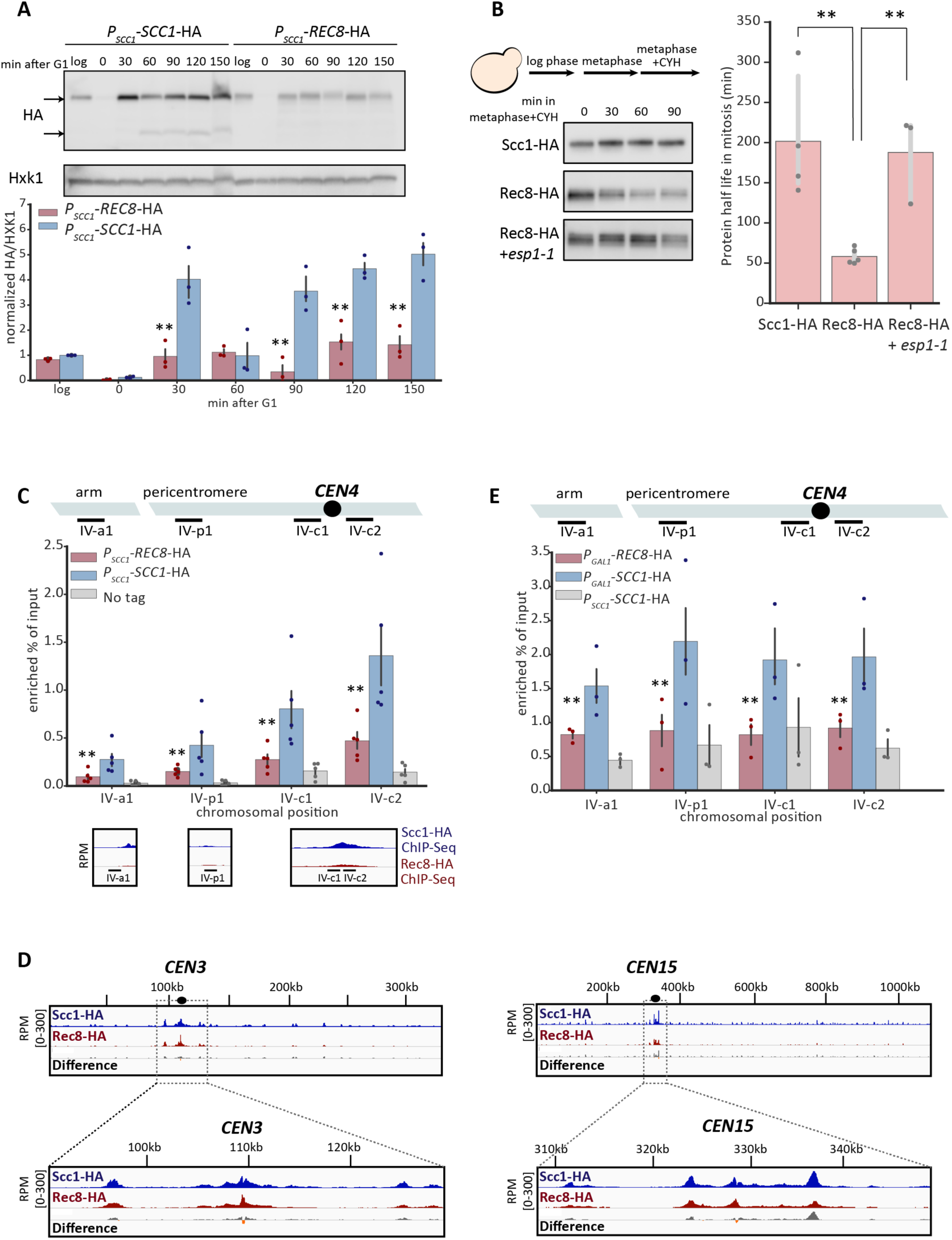
Rec8 is unstable and shows reduced binding to mitotic chromosomes. **(A)** The Rec8 protein level is lower than Scc1 in mitosis. Both *SCC1* and *REC8* were expressed from the *SCC1* promoter and fused to a triple hemagluttinin tag (3xHA) at their C-termini in a strain that also carried *P_GAL1_*-*SCC1*. To follow kleisin protein levels in a single mitotic cycle, expression from the *GAL1* promoter was repressed and cells were released from a G1 arrest and allowed to proceed through the cell cycle as in Fig. 1D. Cells were collected at the indicated timepoints and cell extracts were obtained by alkaline lysis prior to analysis by Western Blotting. Hxk1 was used as a loading control. In the bar graph, the upper arrow marks the size of full-length protein and the bottom arrow marks the size of cleavage product. The three colored points represent the values of three biological replicates and the dark gray bar represents one standard deviation on each side of the mean of these measurements. The statistical significance between data from the Rec8-expressing strain and wild type was calculated by two-tailed Student *t* test, ** *p* < 0.01. **(B)** The instability of Rec8 in mitosis depends on separase activity. Cells were grown to log phase in YPD at 30°C and held in mitosis by addition of benomyl. To check protein stability, cycloheximide was added to the cultures to inhibit protein synthesis. Cells were collected every 30 minutes to examine protein level by Western Blotting. Both Scc1 and Rec8 were detected by an anti-HA antibody. The darker gray points represent the values of three or four biological replicates and the thinner gray bar represents one standard deviation on each side of the mean of these measurements. The half-life of Rec8 is increased in cells expressing a temperature-sensitive mutant of Esp1 (*esp1-1*). (two-tailed Student *t* test, ** *p* < 0.01) **(C)** The level of chromosome-bound Rec8 is lower than that of Scc1 in mitosis. Strains were released from a G1 arrest, proceeded synchronously through one cell cycle and then were arrested in mitosis in YPD containing benomyl. The chromosome-bound kleisin proteins was immunoprecipitated using an anti-HA antibody. Chromatin lysates were prepared from wild type, a Rec8-expressing strain, and a strain without HA tag as negative control. The level of chromosome bound kleisin at the known cohesin binding sites was measured by the amount of DNA that associated with the immunoprecipitated kleisin. DNA was measured by qPCR and expressed as the fraction of material compared to the total chromatin lysate (shown in the y-axis). Four genomic loci on chromosome IV are shown. The colored points represent the values of five biological replicates and the dark gray bar represents one standard deviation on each side of the mean of these measurements. The statistical significance between data from the Rec8-expressing strain and wild type was calculated by two-tailed Student *t* test, ** *p* < 0.01. The bottom panel shows the ChIP-Seq data of Scc1 and Rec8 at the corresponding cohesin binding sites, under the same conditions. **(D)** ChIP-Seq analysis of Scc1- and Rec8-binding in mitosis. Sample preparation and immunoprecipitation were done as in Fig. 2C. The immunoprecipitated DNA bound by Scc1 or Rec8 were examined by whole genome sequencing. The amount of immunoprecipitated DNA is expressed as reads per million (RPM) calibrated to the reference, *S. pombe* genomic DNA immunoprecipitated by the Scc1 ortholog, Rad21. The calibrated signal of ChIP-seq data (relative to a control *S. pombe* sample) representing the degree of enrichment of kleisin (Scc1 in blue and Rec8 in red) and the difference in enrichment between the two kleisins (gray: Scc1’s signal is more than Rec8’s; orange: Rec8’s signal is more than Scc1’s) is visualized by the Integrated Genome Viewer (Robinson et al., 2011). Chromosome III and XV are shown as examples of chromosomes with different sizes and different degrees of peri-centromeric Rec8 binding, with an expanded view of 20kb DNA on each side of a centromere in the bottom panel. **(E)** Rec8 loads poorly on G1 chromosomes compared to Scc1. To overexpress kleisins in G1, the *P_GAL1_*-*SCC1*-HA and *P_GAL1_*-*REC8*-HA strains were arrested in YEP containing 2% galactose and alpha-factor. Chromatin immunoprecipitation and qPCR were performed as described in Fig. 2B. The darker points represent the values of three biological replicates and the darker gray bar represents one standard deviation on each side of the mean of these measurements. The statistical significance between data from the Rec8-expressing strain and wild type was calculated by two-tailed Student *t* test, ** *p* < 0.01.

Finally, we compared the binding of Rec8 and Scc1 to chromosomes by chromatin immunoprecipitation (ChIP). In mitotically-arrested cells, immunoprecipitating Rec8 brought down less DNA at canonical cohesin binding sites than Scc1 (Fig. 2C). This was not only true for individual sites, but was also observed genome-wide. Calibrated ChIP-Seq revealed reduced levels of chromosomal Rec8 compared to Scc1 at peri-centromeres, where cohesin is most enriched, across all the sixteen chromosomes (Fig. 2D and Fig. S4A). Among all the chromosomes, the enrichment of Rec8 specifically at core centromeres is higher compared to that of Scc1, but lower in the flanking peri-centromeres (Fig. 2D and Fig. S4B). This suggests that Rec8-containing cohesin is less efficient than Scc1 in translocating from its loading site at centromere. The reduced overall binding of Rec8 on mitotic chromosomes might be partially explained by the lower Rec8 level in mitosis (Fig. S5A). To test this idea, we asked whether equivalent amounts of ectopically produced Rec8 and Scc1 can be loaded on chromosomes in G1. Although Scc1 is not normally expressed until the onset of S phase, ectopically expressing Scc1 in G1 allows cohesin to be loaded on chromosomes (Fernius et al., 2013). We expressed Rec8 or Scc1 from the *GAL1* promoter in G1-arrested cells and measured their chromatin association by ChIP-qPCR. Although the levels of ectopically produced Rec8 was slightly elevated compared to that of Scc1 (Fig. S5B), it showed reduced accumulation on canonical cohesin binding sites (Fig. 2E). We conclude that Rec8 both is less stable and, independently, associates less well with chromosomes compared to Scc1. We suggest that these molecular defects lead to defective sister cohesion, errors in chromosome segregation, and slower passage through mitosis, leading to the production of dead and aneuploid cells, thereby reducing the fitness of Rec8-expressing cells.

### Experimental evolution increases the fitness of Rec8-expressing strains

To study how cells adapt to a protein that performs an essential function poorly, we asked if experimental evolution would allow Rec8-expressing cells to acquire mutations that would improve their fitness; these mutations could occur either in *REC8* or elsewhere in the yeast genome. We constructed fifteen ancestral clones, each containing a deletion of the chromosomal *SCC1* gene and a centromeric plasmid expressing *REC8* from the *SCC1* promoter (*P_SCC1_-REC8*). Ancestral clones were inoculated into rich medium at 30°C and each culture was diluted 6000-fold into fresh medium once it reached saturation. This process was repeated, freezing samples every 125 generations, until the populations reached 1750 generations (Fig. 3A). At generation 375, the fitness of all the evolved populations had increased by 20-30% relative to the Rec8-expressing ancestor. At the end of the experiment, the fitness of the evolved populations was 30-80% greater than that of the ancestor and the fitness of two evolved populations (P13 and P15) was similar to the wild-type strain (Fig. 3B and Fig. S6).

**Figure 3.**
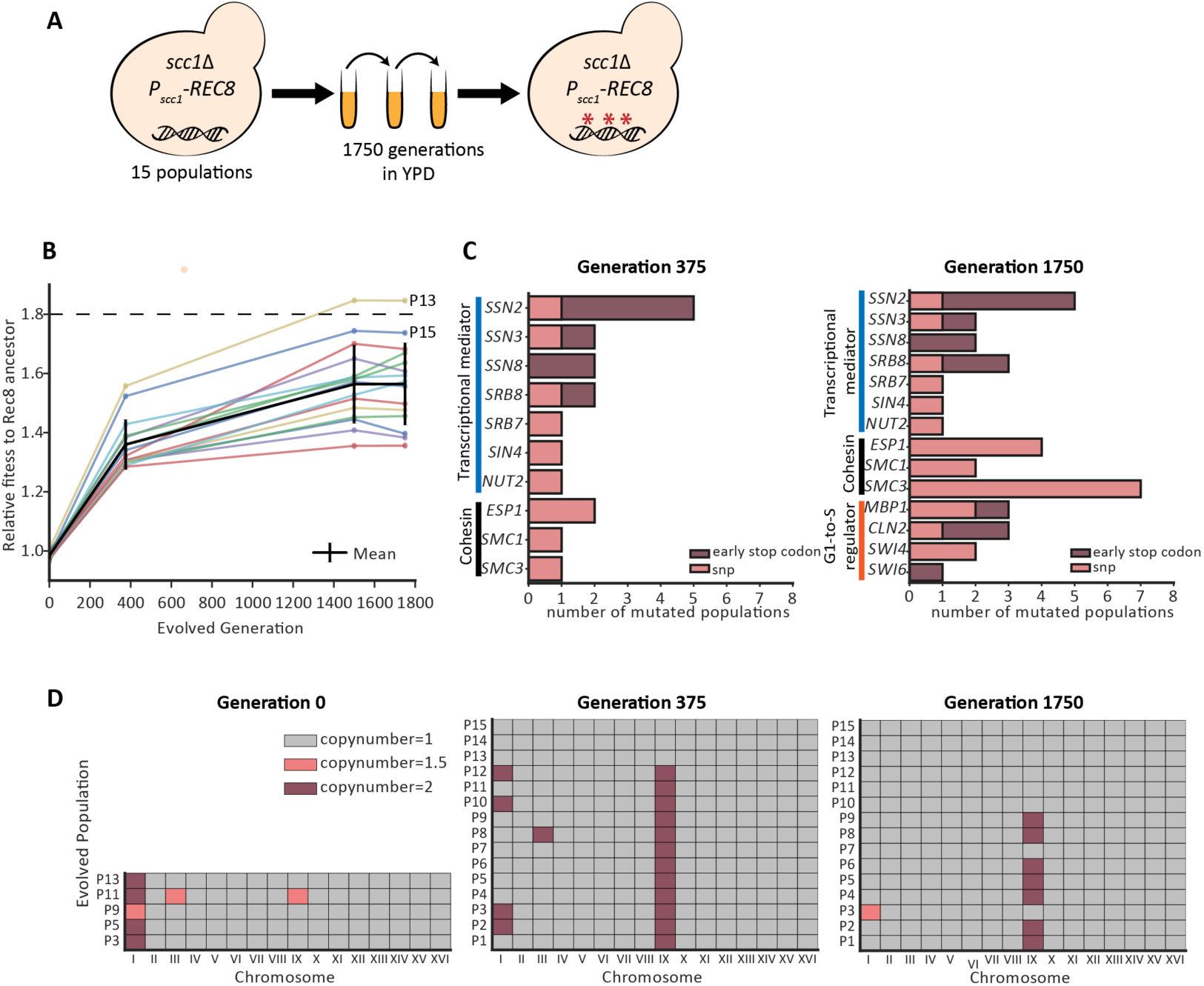
Experimental evolution improves the fitness of Rec8-expressing populations. **(A)** Schematic of the experimental evolution of fifteen independent populations forced to use Rec8 in mitosis for 1750 generations **(B)** The fitness of all the evolved populations increases during evolution. The relative fitness of each evolved population at generation 375 and generation 1750 was measured by competing them against a fluorescently-labeled ancestor in YPD. Changes in the relative fitness of individual evolved population during evolution is shown as individual colored line. The average fitness of all 15 evolved populations is shown as a black line. The fitness of wild type relative to the Rec8-expressing ancestor is indicated as a black dashed line. Two evolved populations showing fitness near wild type are labeled (P13 and P15). **(C)** Summary of functional modules that had acquired fixed mutations in more than six populations at generation 1750. The x-axis shows the number of populations that acquire a mutation in any specified gene at generation 375 and 1750. The y-axis shows mutated genes grouped by their functions: genes involved in the transcriptional mediator complex in blue, cohesin related genes in black, the G1-to-S cell cycle regulators in orange. Mutations causing early stop codon are shown in dark red and single nucleotide changes are shown in pink. **(D)** Summary of changes in chromosomal copy number of all fifteen evolved populations. The copy number of each chromosome was calculated by normalizing the read depth spanning each chromosome to the median read depth over the entire genome. The results of five ancestral clones and fifteen evolved populations at generation 375 and 1750 are shown here: gray marks one copy, dark red marks two copies, and pink marks 1.5 copies, suggesting part of population were disomic.

To identify adaptive mutations, we sequenced the genomes of five ancestral clones and pooled genomes of fifteen evolved populations at generation 375 and generation 1750 (Sup File 1). We focused on non-synonymous mutations that were present at a frequency ≥90% in any evolved population. The evolved populations had an average of nine mutations at generation 375 and 17 mutations at generation 1750 that met this criterion. We did not find any mutations in *REC8*, either in the coding sequence or the DNA 500 bp upstream and downstream of the ORF, but we did find multiple mutations in three functional modules: the transcriptional mediator complex, cohesin and its regulators, and regulators of cell cycle progression from G1 to S phase. At generation 375, fourteen out of fifteen evolved populations had a mutation in the transcriptional mediator complex, and four populations had mutations in the other two cohesin subunits, *SMC1* and *SMC3*, or separase, *ESP1* (Fig. S6 and Table S1). At generation 1750, the early mediator mutations were still fixed, one population had acquired a mutation in a second mediator subunit (*SRB8*) and one population still lacked a mediator mutation (Fig. S6 and Table S2). Twelve out of the fifteen mediator mutations targeted the Cdk8 complex, a regulatory module of mediator, and nine of these twelve mutations produced early stop codons (Fig. 3C). Mutations in cohesin-related genes were common at generation 1750: *SMC3, SMC1*, and *ESP1* were mutated in seven, two, and four evolved populations respectively (Fig. 3C). Seven populations had mutations in one of these genes, three populations had mutations in two genes and five populations had not acquired mutations in any cohesin-related gene by generation 1750. Four genes (*MBP1*, *CLN2*, *SWI6*, and *SWI4*) controlling the cell cycle transition from G1 to S were mutated in a total of six populations at generation 1750 (Fig. 3C). In summary, nine out of the fifteen evolved populations acquired mutations both in the mediator complex and cohesin-related genes (Fig. S6). Only the three fittest populations had mutations in all three classes (cohesin-related, mediator, and G1-to-S regulators) and the fitness of two of these populations, P13 and P15, approached that of wild type (Fig. S6).

In addition to point mutations, many evolved populations were aneuploid (Fig. 3D). The five ancestral clones we sequenced had an extra copy of chromosome I, the smallest chromosome in budding yeast. We think this reflects a combination of three factors, a very high frequency of chromosome mis-segregation in the ancestral Rec8-expressing cells, preferential mis-segregation of smaller chromosomes, and the small fitness cost of an extra copy of chromosome I (Torres et al., 2007). At generation 375, twelve populations had independently gained an extra copy of chromosome IX, and five also had an extra copy of chromosome I or chromosome III. The other three were true haploids, with one having lost the extra copy of chromosome I that was present in its ancestor. At generation 1750, seven populations retained two copies of chromosome IX while the rest had become true haploids. Disomy for chromosome IX causes a slight fitness cost in wild type (Torres et al., 2007), but in our evolution experiment, the prevalence and persistence of chromosome IX disomes suggests that an extra copy of chromosome IX in Rec8-expressing cells is adaptive.

The genetic alterations we found are specific to yeast cells adapting to expressing Rec8 rather than Scc1. Mutations in transcriptional mediator, cohesin-related genes, and the G1-to-S regulators have not been seen at frequencies that suggest they are adaptive in previous experimental evolution studies in *S. cerevisiae* (Jerison et al., 2017; Kryazhimskiy et al., 2014; Laan et al., 2015; Lang et al., 2013). In evolution experiments that improve the growth of haploid yeast in rich medium, the most frequent ploidy change seen is diploidization (Gerstein et al., 2006; Kryazhimskiy et al., 2014), instead of gaining an extra copy of a specific chromosome, which has been seen for cells adapting to the absence of myosin (Rancati et al., 2008) and growth at high temperature (Yona et al., 2012).

### Reconstruction confirms that candidate mutations are adaptive

We tested the effect of putative causative mutations by engineering them, individually, into the Rec8-expressing ancestor and examining the fitness and phenotypes of the resulting strains. We focused on four groups of genetic changes: mutations in transcriptional mediator, cohesin components, regulators of the G1-to-S transition, and an extra copy of chromosome IX. Mutations in the transcriptional mediator complex primarily targeted the Cdk8 complex: of its four components, *SSN2* was mutated five times, *SSN3* and *SSN8* were each mutated twice, and *SRB8* was mutated three times. Nine out of these twelve mutations led to early stop codons, suggesting that these mutations inactivate the module’s function. The mediator complex links the basic transcriptional machinery with transcription factors and controls various events in transcription, including transcriptional initiation, pausing, elongation, and the organization of chromatin structure (Allen and Taatjes, 2015). The Cdk8 kinase module of mediator can positively or negatively regulate transcription (Nemet et al., 2014). We reconstructed mutations in three subunits (*SSN2*, *SSN3*, and *SSN8*) of the Cdk8 module. Each increased the fitness of the ancestor by 8-25% (Fig. 4A). Deleting the above genes also increased the fitness of the ancestor (Fig. 4A), strongly suggesting that these evolved mutations are loss-of-function mutations. Of the mutations targeting cell cycle regulators, one of three mutations in *MBP1* and two of three mutations in *CLN2* caused early stop codons. We therefore mimicked the effect of these mutations by deleting the corresponding gene: *mbp1*Δ and *cln2*Δ increased the fitness of the ancestor by 20% and 25%, respectively (Fig. 4B).

**Figure 4.**
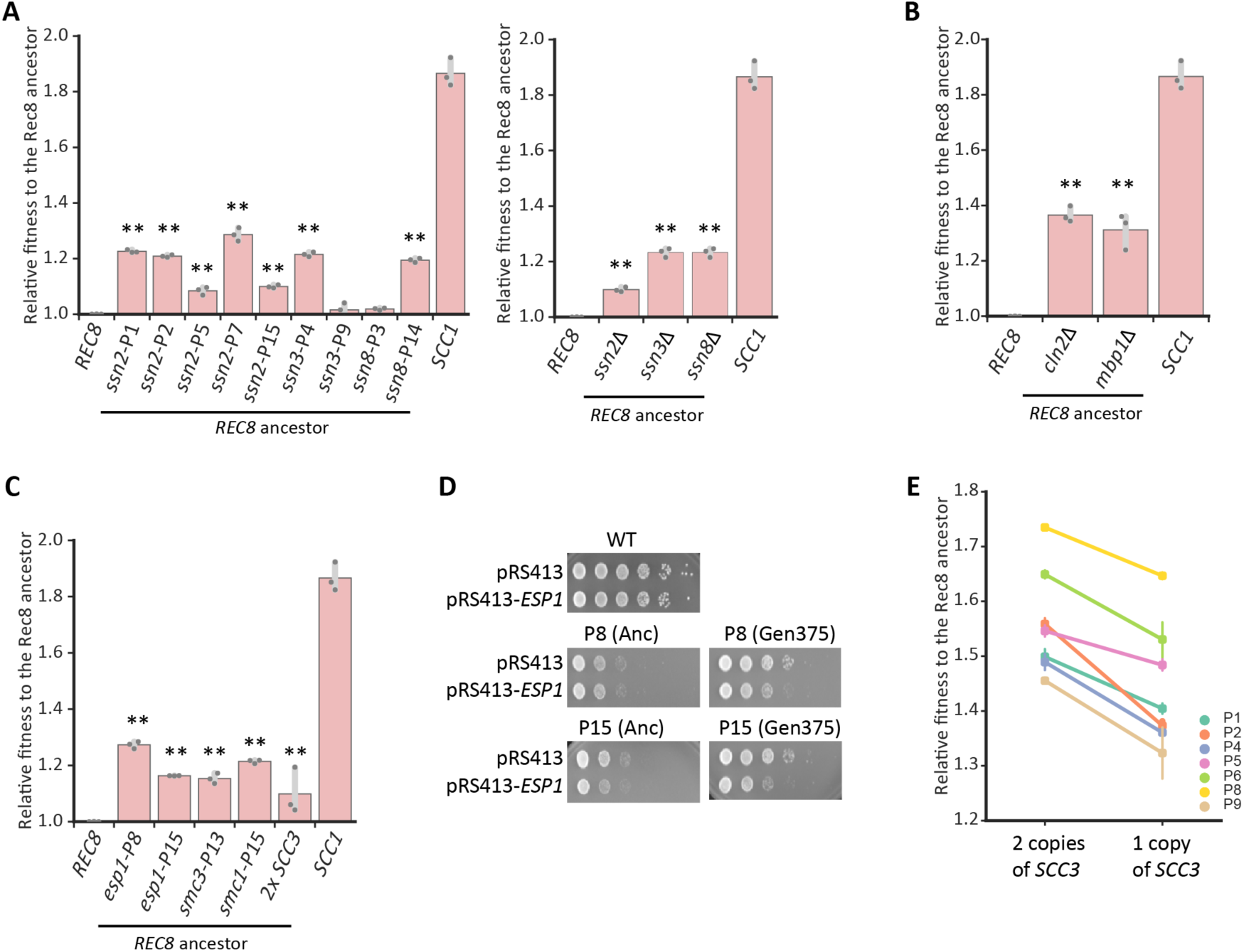
Reconstructing individual evolved mutations increases the fitness of the Rec8-expressing ancestor. **(A)** The effect of single evolved mutation and deletion of genes encoding subunits of the Cdk8 complex on fitness of the Rec8-expressing ancestor. **(B)** The effect of deleting *CLN2* and *MBP1* on fitness of the Rec8-expressing ancestor. **(C)** The effect of single evolved mutations in genes that encode other cohesin components or separase and an extra copy of *SCC3* on fitness of the Rec8-expressing ancestor. In **4A-4C**, each single evolved mutation was reconstructed in the Rec8 ancestral strain used in the evolution experiment. The relative fitness of reconstructed strains to the ancestor was measured by competing it against a fluorescently-labeled Rec8 ancestor. The darker gray points represent the values of three biological replicates and the thinner gray bar represents one standard deviation on each side of the mean of these measurements. The fitness of the wild-type strain, labeled as *SCC1*, is shown in each panel. The statistical significance between data from the Rec8-expressing strain and each mutation-reconstructed strain was calculated by two-tailed Student *t* test, ** *p* < 0.01. **(D)** *esp1* evolved mutations (*esp1-P8* and *esp1-P15*) are hypomorphic. A *CEN* plasmid carrying *ESP1* was transformed into a wild-type strain, two ancestors (Anc), and two evolved populations that had acquired *esp1* mutations (P8 and P15) at generation 375. Cells were subjected to ten-fold serial dilutions and spotted on YPD plates to assay growth. Cells transformed with an empty plasmid (pRS413) served as control. **(E)** The effect of deleting one copy of *SCC3* of fitness of the evolved populations with disomic chromosome IX at generation 1750.

The mutations in cohesin-related genes affect essential genes and are thus unlikely to eliminate the function of these genes. Individual mutations in *ESP1, SMC1,* and *SMC3* increased the fitness of the Rec8-expressing ancestor by 15-31%, 14%, and 21% respectively (Fig. 4C). Our finding that Rec8 is sensitive to separase activity in mitosis (Fig. 2B) raised the possibility that evolved *esp1* mutations are hypomorphic alleles that weaken separase activity. We tested this hypothesis by expressing an extra wild-type copy of *ESP1* in two evolved populations carrying *esp1* mutations and their ancestors. As predicted, the extra copy of *ESP1* reduced the growth of these two evolved populations but not their ancestors (Fig. 4D), suggesting that these evolved *esp1* mutations are hypomorphic. Consistent with this hypothesis, compromising separase activity by using a known temperature-sensitive mutation, *esp1-1* (Ho et al., 2015), increased the growth of the Rec8-expressing ancestor at the permissive temperature (Fig. S7).

The prevalence of chromosome IX disomy in our evolved populations suggested that two copies of chromosome IX confer a selective advantage on Rec8-expressing strains. Aneuploidy has been adaptive in several evolution experiments by increasing the copy number of a specific gene (Mangado et al., 2018; Rancati et al., 2008; Sunshine et al., 2015; Voordeckers et al., 2015). Chromosome IX encodes a candidate gene, *SCC3*, whose protein product promotes cohesin association with chromosomes (Roig et al., 2014) by interacting with the cohesin loading complex (Orgil et al., 2015) and Scc1 (Li et al., 2018). We asked if an extra copy of *SCC3*, in the absence of the other genes on chromosome IX, could increase the fitness of the Rec8-expressing ancestor. We integrated an extra copy of *SCC3* in the ancestor and found that this manipulation increased its fitness by 10% (Fig. 4C), demonstrating that an extra copy of *SCC3* is sufficient to increase fitness. To test if an extra copy of *SCC3* is also necessary for increasing fitness, we deleted one copy of *SCC3* in clones from seven evolved populations that carried two copies of chromosome IX at generation 1750, reducing their fitness by 8 to 28% (Fig. 4E). We conclude that an extra copy of *SCC3* explains much of the selective advantage of carrying an extra copy of chromosome IX.

### Adaptive genetic changes restore sister chromosome cohesion

Do the adaptive mutations in Rec8-expressing strains increase fitness by improving sister cohesion? We engineered individual mutations into a Rec8-expressing strain containing a GFP-labeled chromosome V and *P_GAL1_-SCC1* and examined sister chromosome cohesion after acute depletion of Scc1. Individually deleting three subunits in the Cdk8 complex partially rescued the sister chromosome cohesion defect in cells where Rec8 was the only α-kleisin present (Fig. 5A and Fig. S8). Amongst these genes, deleting *SSN3*, the kinase subunit of the Cdk8 complex, produced the greatest improvement in sister cohesion, comparable to the effect of the adaptive mutations in cohesin-related genes (*SMC1*, *SMC3*, or *ESP1*) or deleting the two genes that promote exit from G1, *CLN2* and *MBP1* (Fig. 5A). An extra copy of *SCC3*, whose effect on fitness mimicked the chromosome IX disome, slightly improved sister cohesion (Fig. 5A). Each evolved mutation also improved the accuracy of chromosome segregation in Rec8-expressing cells with cohesin-related mutations having stronger effects than mediator mutations (Fig. S9). We conclude that mutations in transcriptional mediator, other cohesin components and separase, and cell cycle regulators can improve sister chromosome cohesion in Rec8-expressing cells.

**Figure 5.**
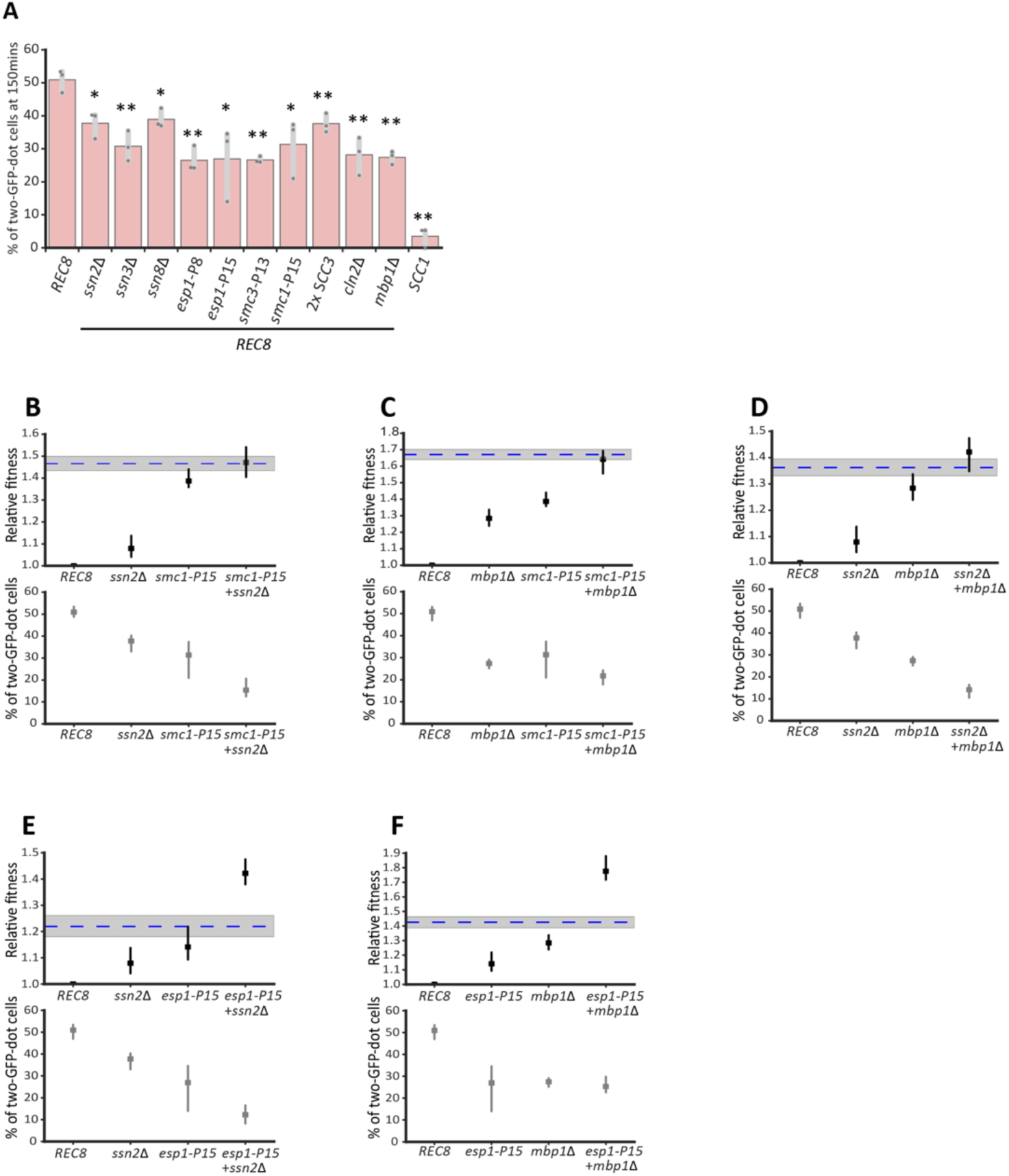
Adaptive genetic changes improve sister chromosome cohesion in Rec8-expressing cells. **(A)** Individual adaptive genetic changes partially improve sister chromosome cohesion Deletions of genes in the Cdk8 complex, adaptive mutations in cohesin and its regulator, two copies of *SCC3*, and deletions of genes that regulate G1-to-S transition were reconstructed individually in the strain used for assaying sister cohesion. Cells were prepared as in Fig. 1C, and the percentage of cells with two GFP-dots in populations arrested in mitosis (150mins after releasing from G1) is shown. The darker gray points represent the values of three biological replicates and the thinner gray bar represents one standard deviation on each side of the mean of these measurements. The statistical significance between data from the Rec8-expressing strain and each mutation-reconstructed strain was calculated by two-tailed Student *t* test, * *p* < 0.05, ** *p* < 0.01. **(B-F)** Relative fitness and sister chromosome cohesion of double mutants are shown: *ssn2*Δ and *smc1-P15* (B), *mbp1*Δ and *smc1-P15* (C), *ssn2*Δ and *mbp1*Δ (D), *ssn2*Δ and *esp1-P15* (E), *mbp1*Δ and *esp1-P15* (F). The blue dashed line represents the expected fitness if two mutations contribute additively, and the shaded region represents the standard error of that expectation

We investigated the interactions between adaptive mutations in different functional modules. The fitness of the evolved population P15 approached that of wild type at generation 1750 and it had acquired mutations in four genes (*ssn2*, *esp1*, *smc1*, and *mbp1*) representing effects on mediator, separase, cohesin, and the G1-to-S transition. We investigated the interaction between these four mutations. To examine fitness and sister chromosome cohesion, we constructed double mutants in the strain carrying a GFP-labeled chromosome V and *P_GAL1_*-*SCC1*. For fitness, we saw two types of interactions: double mutations between any of *ssn2*Δ, *mbp1*Δ, and *smc1-P15* had a fitness that was indistinguishable from the sum of the effects of the individual mutations (Fig. 5B-D), whereas the *esp1-P15 ssn2*Δ and *esp1-P15 mbp1*Δ double mutants were substantially fitter than the sum of the fitness increases in the individual mutants (Fig. 5E and 5F). For sister chromosome cohesion, all the double mutants had smaller defects in sister chromosome cohesion than either single mutant with the exception of the *mbp1*Δ *esp1-P15* and *mbp1*Δ *smc1-P15* double mutants (Fig. 5C and 5F), whose level of sister chromosome cohesion was either indistinguishable from or only slightly above that of the single mutants. This result suggests that *mbp1*Δ may have additional effects on fitness that are not mediated by improving sister chromosome cohesion. Overall, the interactions between mutations in different modules are additive or positively synergistic at the level of fitness and more complex at the level of sister cohesion.

We asked if the adaptive mutations altered the abundance of Rec8. We measured the Rec8 protein level in mitosis in seven strains, each containing an adaptive mutation in a different gene that appeared during our evolution experiment and was shown to increase the fitness of Rec8-expressing cells (Fig. S10). None of the mutations changed the level of Rec8, demonstrating that these adaptive mutations improve sister cohesion not by changing the amount of Rec8.

### Adaptive genetic changes slow down S phase and improve sister cohesion

Cell cycle progression profiles showed that the ancestral Rec8-expressing strain had shorter S phase than that of wild type (Fig. 1D). Since the linkage between sister chromosomes is established in S phase (Uhlmann and Nasmyth, 1998) and all the adaptive mutations improved sister chromosome cohesion in the Rec8-expressing strain, we asked if these mutations also affected the dynamics of genome replication. By tracking cell cycle progression after release from a G1 arrest, we found deletions of three subunits in the Cdk8 complex and mutations in *SMC1, SMC3,* or *ESP1* slowed S phase of the Rec8-expressing strain (Fig. 6A). We quantified the fraction of cells in S phase 30 minutes after release from a G1 arrest: 39% of wild-type cells were in S phase, whereas 57% of the Rec8-expressing cells were in S phase. In the Rec8-expressing strain, deleting genes encoding the subunits of the Cdk8 complex decreased the fraction of cells in S phase to between 50 and 33% and mutations in *SMC1, SMC3,* and *ESP1* decreased this fraction to between 40% and 17% (Fig. 6B and Fig. S11). The budding index of these reconstructed strains are comparable to those of both the wild-type and Rec8-expressing strains (Fig. S12), confirming that none of the mutations affect the timing of Start after release from a G1 arrest.

**Figure 6.**
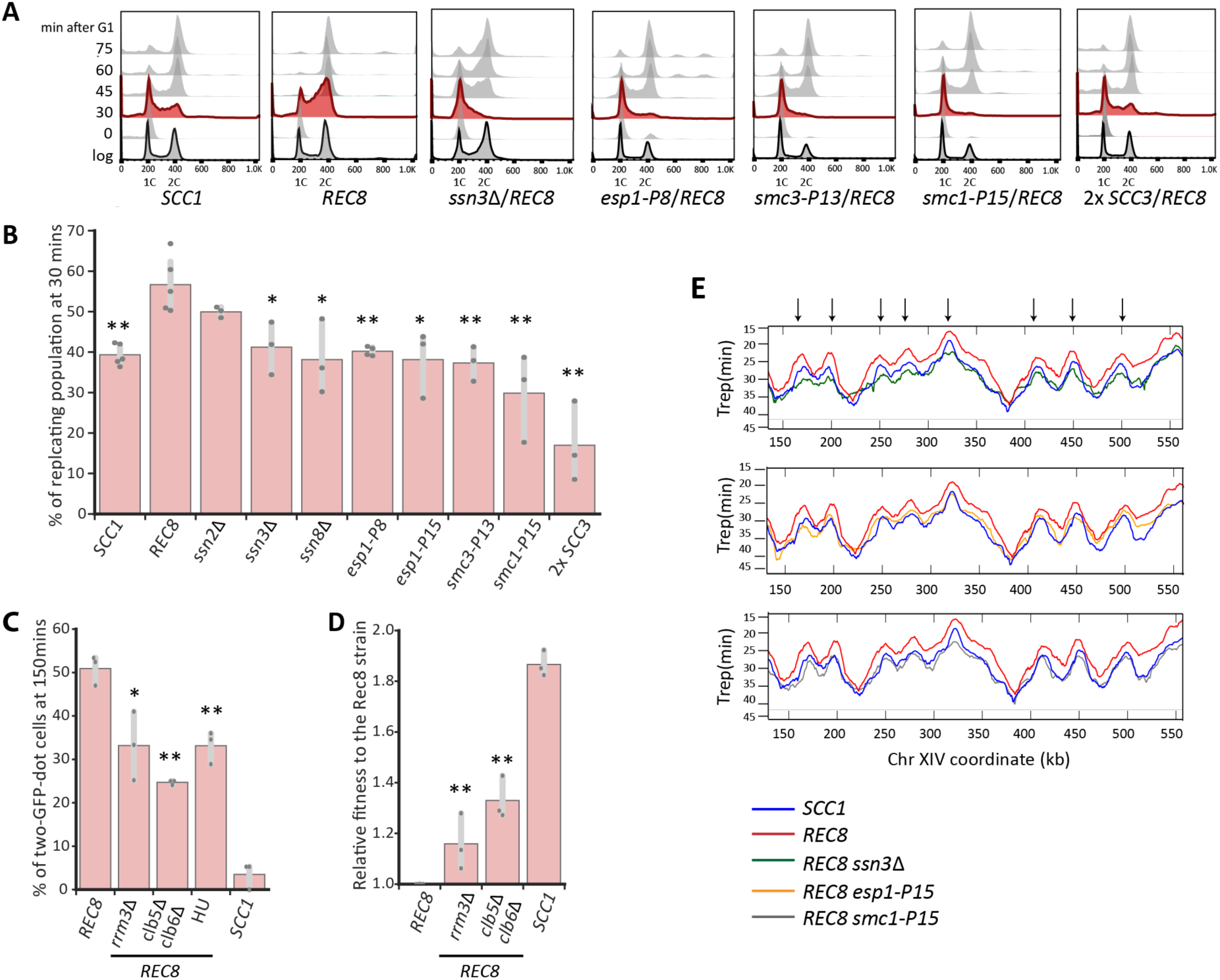
Slowing down genome replication partially improves sister chromosome cohesion in Rec8-expressing cells. **(A)** Cell cycle profiles of wild type strain, the Rec8-expressing strain, and the Rec8-expressing strains carrying a single reconstructed mutation. Cells were released from the G1 arrest as described in Fig. 1D. Flow cytometry profiles are shown at the indicated times and profiles at 30 minutes are labeled in red. **(B)** Quantitation of the fraction of replicating cells in strains carrying a single reconstructed mutation at 30 minutes after release from a G1 arrest. The replicating subpopulation was measured as the fraction of the population between the G1 peak and the G2/M peak. The darker gray points represent the values of three biological replicates and the thinner gray bar represents one standard deviation on each side of the mean of these measurements. The statistical significance between data from the Rec8-expressing strain and each mutation-reconstructed strain was calculated by two-tailed Student *t* test, * *p* < 0.05, ** *p* < 0.01. **(C)** Genetically and chemically perturbing genome replication improves sister chromosome cohesion. The Rec8-expressing *rrm3*Δ and *clb5Δ clb6*Δ strains were assayed for sister chromosome cohesion as described in Fig. 1C. 12.5mM hydroxyurea was added in YPD as Rec8-expressing cells entered the cell cycle. The percentages of cells with two GFP-dot in mitotically-arrested populations (150mins after G1) are shown. The darker gray points represent the values of three biological replicates and the thinner gray bar represents one standard deviation on each side of the mean of these measurements. The statistical significance between data from the Rec8-expressing strain and each mutant strain was calculated by two-tailed Student *t* test, * *p* < 0.05, ** *p* < 0.01. **(D)** The effect of *rrm3*Δ and *clb5Δ clb6*Δ on the fitness of the Rec8-expressing strain. The darker gray points represent the values of three biological replicates and the thinner gray bar represents one standard deviation on each side of the mean of these measurements. The statistical significance between data from the Rec8-expressing strain and each mutant strain was calculated by two-tailed Student *t* test, ** *p* < 0.01. **(E)** The replication profiles of wild type, the Rec8-expressing strain, and the Rec8-expressing strains with a single reconstructed mutation (*ssn3*Δ, *esp1-P15*, or *smc1-P15*). Replication dynamics is expressed as T_rep_ (shown in the y-axis), the time at which 50% of cells in a population complete replication at a given genomic locus. The mean replication profile of two experiments on one part of chromosome XIV is shown. The replication profile of each strain is color-coded. An arrowhead represents a fired replication origin. We confirmed that the different strains exited from G1 at the same time by monitoring their budding index over time (Fig. S15).

We asked how Rec8 altered genome replication and how individual adaptive mutations restored the tempo of genome replication to that of wild-type cells. We constructed a whole genome replication profile by genome sequencing multiple time points of a synchronized yeast population proceeding through S phase (Saayman et al., 2018). By analyzing changes in read depth during S phase, we calculated T_rep_, the time at which 50% of cells in a population complete replication at a given genomic locus. The profile of T_rep_ across the yeast genome reveals the dynamics of replication: the peaks mark points at which replication initiates, namely a fired replication origin, and the slopes show the speed of the replication forks that move away from the origins. We compared the replication profiles of wild type, *scc1*Δ, Rec8-expressing, and reconstructed strains that express Rec8 and carry a single adaptive mutation in one of three genes: *ssn3*Δ, *esp1-P15*, or *smc1-P15*. Compared to the wild-type, the Rec8-expressing strain fired many but not all replication origins earlier, and on average an origin fired four minutes earlier in the Rec8-expressing cells. In contrast, the temporal order of origin firing and the speed of replication forks were similar to those in wild type. Because origins that fire earlier are less likely to be inactivated by the nearby origins and therefore replicate DNA more efficiently (Bell and Labib, 2016), this result is consistent with the earlier S-phase of the Rec8-expressing strain. The replication profile of *scc1*Δ strain was indistinguishable from that of the wild-type (Fig. S13). Three adaptive mutations we examined delayed origin firing to various degrees: *ssn3*Δ or *esp1-P15* almost restored the pattern of origin firing to that of the wild-type. *smc1-P15* made the firing of many origins later than those of wild type (Fig. 6E and Fig. S14). Overall, the replication profiles of these reconstructed strains are more similar to the genome-wide pattern of wild type rather than that of the Rec8-expressing strain. We concluded that expressing Rec8 in mitosis advances the timing of origin firing and therefore Rec8-expressing cells begin and finish genome replication earlier than wild type. Mutations in genes encoding the Cdk8 complex, cohesin and separase slow down the S phase of the Rec8-expressing strain by delaying origin firing. In Scc1-expressing cells, deleting the genes encoding the subunits of the Cdk8 complex extended S phase and led to an 8-11% fitness reduction (Fig. S16), demonstrating that transcriptional mediator affects genome replication, even in the absence of Rec8, through an uncharacterized mechanism.

The correlation between slower genome replication and improved sister chromosome cohesion suggests that the dynamics of genome replication affect cohesion. Cohesin must be loaded onto chromosomes prior to, or concomitant with the passage of the replication fork to be converted into functional cohesion (Uhlmann and Nasmyth, 1998) and delaying origin firing promotes the establishment of cohesive linkages near centromeres in a kinetochore mutant defective in cohesin accumulation at centromeres (Fernius and Marston, 2009). Based on these observations, we hypothesized that slowing down genome replication would improve Rec8-dependent sister chromosome cohesion. To test this idea, we asked if manipulations that slow genome replication improve sister chromosome cohesion and the fitness of the Rec8-expressing strain, potentially by allowing more time for cohesin to load prior to replication fork passage. We found that both decreasing origin firing and slowing the movement of replication forks improved sister cohesion. Removing the two S phase cyclins, Clb5 and Clb6, which delay replication origin firing (Donaldson et al., 1998; Schwob and Nasmyth, 1993), or reducing the speed of replication forks by removing Rrm3, a helicase involved in DNA replication (Azvolinsky et al., 2006), halved the sister cohesion defect in Rec8-expressing cells (Fig. 6C) and increased their fitness by 31% (*clb5*Δ *clb6*Δ) and 24% (*rrm3*Δ) (Fig. 6D). Slowing genome replication with hydroxyurea (HU), which lowers the concentration of deoxyribonucleotide triphosphates (dNTPs), also improved sister cohesion and fitness (Fig. 6C and Fig. S17). We infer that in Rec8-expressing cells, the Cdk8 and cohesin-related mutations exert at least part of their effects by slowing genome replication and thus improving sister cohesion (Fig. 7).

**Figure 7.**
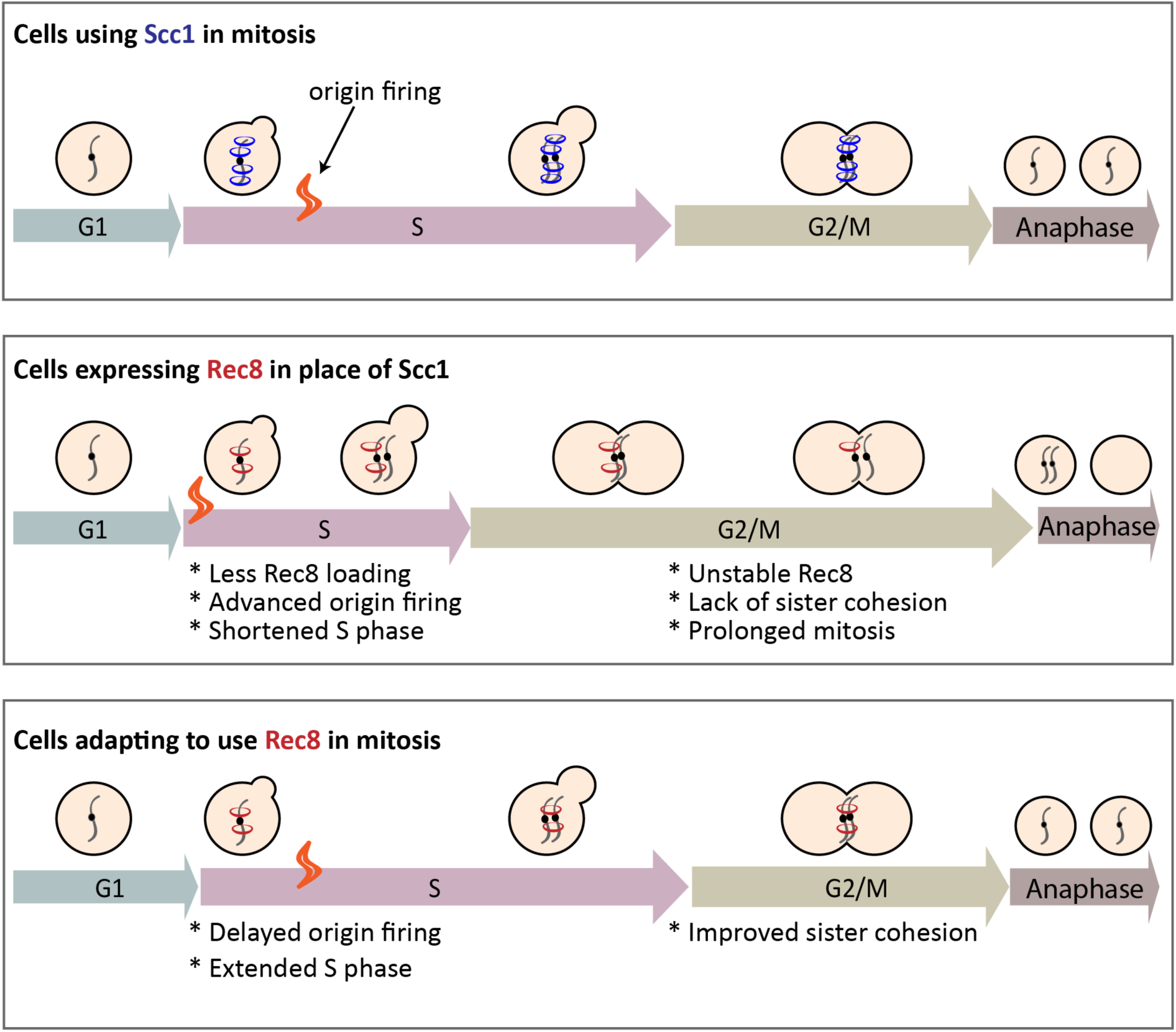
Summary of the mechanism that allows budding yeast to use the meiotic kleisin, Rec8, for mitosis. Yeast cells expressing Rec8 in place of Scc1 cannot build robust cohesion to hold sister chromosomes together before anaphase due to the weak association of cohesin with chromosomes and Rec8 protein instability. Rec8-expressing cells induce earlier firing of replication origins compared to wild type does and exhibits shortened S phase. After experimental evolution, adaptive mutations in different functional modules delay origin firing and improve sister chromosome cohesion of cells that are forced to use Rec8 in mitosis, potentially by allowing more time for Rec8-cohesin to load onto chromosomes prior to passage of the replication fork.

## Discussion

We used experimental evolution to study how cells adapt to the demand that a protein performs an altered function. Budding yeast adapt to use the meiotic kleisin, Rec8, which normally functions in meiosis, to maintain the sister chromosome linkage required for accurate mitotic chromosome segregation. Whole genome sequencing of the adapted populations failed to reveal mutations in *REC8* but identified adaptive mutations in three functional modules: the transcriptional mediator complex, cohesin structure and regulation, and cell cycle regulation. Individually, these mutations slow genome replication, improve sister cohesion, and increase the fitness of the ancestral Rec8-expressing strain. Engineering mutations that delay the firing of replication origins or slow the speed of replication forks into the ancestral Rec8-expressing strain, increased sister chromosome cohesion and fitness, demonstrating a causal link between genome replication and sister chromosome cohesion. Our work suggests that mutations, both in the components and regulators of cohesin and in other proteins, which were not previously implicated in chromosome cohesion, improve the ability of the meiotic kleisin to function in mitosis, despite the passage of a billion years since the divergence between mitotic and meiotic kleisins.

What distinguishes the cellular functions of Scc1 and Rec8? Cells that are forced to use Rec8 in mitosis have multiple defects that account for their reduced fitness. In G1, ectopically-expressed Rec8 associates more weakly with chromosomes than Scc1does. This defect may reduce the ability to productively load Rec8-containing cohesin on chromosomes. In mitotically-arrested cells, Rec8 is less stable than Scc1, binds less well to peri-centromeres and chromosomal arms, and shows increased binding specifically at core centromeres. We suggest that the reduced pericentromeric binding in mitotically-arrested cells destabilizes sister chromosome cohesion. This reduced stability of Rec8 is partially due to separase activity: populations acquired hypomorphic alleles of separase and a temperature-sensitive separase mutant stabilized Rec8 in mitotically-arrested cells. Rec8’s genome-wide binding pattern suggests Rec8 does not associate with chromosomes in the same way to Scc1. Cohesin binding to chromosomes is initiated by cohesin loading, which is either specifically targeted to centromeres and dependent on the Ctf19 kinetochore complex (Hinshaw et al., 2017; Hinshaw et al., 2015), or generally loaded genome-wide. Following loading, cohesin translocates to other parts of chromosomes (Hu et al., 2011; Lengronne et al., 2004). In mitosis, Rec8’s increased binding at centromeres and reduced binding at peri-centromeres suggests that Rec8-containing cohesin can be targeted to centromeres but cannot translocate efficiently to the peri-centromeric borders, where most of Scc1-containing cohesin accumulates to generate linkages between sister chromosome (Paldi et al., 2019). In meiosis, the difference in the stability of the linkage with the two forms of cohesin is reversed: Scc1 near centromeres is not protected from separase activity whereas Rec8 is (Toth et al., 2000). Since the other cohesin subunits and cohesin regulators are present in both the mitotic and meiotic cell cycles, these differences must be due to differential modification of the known cohesin regulators or additional components that interact differently with Rec8 and Scc1. Why can’t Rec8 fully substitute for Scc1 in the mitotic cell cycle? Our results do not distinguish between two possibilities: i) there is a fundamental incompatibility between the functions that kleisins perform in mitosis and meiosis and this incompatibility forced these two kleisin paralogs to diverge from each other, and ii) mutations that impaired Rec8’s ability to support mitosis accumulated by genetic drift rather than selection. In either case, our work reveals the power of a variety of adaptive mutations, affecting diverse modules, to alter the function that a protein performs.

What accounts for the genes that acquired adaptive mutations and the order in which mutations appear? We argue that the answer is a combination of the benefit conferred by mutations in a gene and the target size for these beneficial mutations. The mutations in the transcriptional mediator complex and genes regulating G1-to-S transition are likely to be strong loss-of-function mutations: many of the mutations are nonsense mutations and gene deletions mimic the effect of the evolved mutations. Two arguments suggest that the mutations in cohesin and its regulators are different: these are essential genes and their mutations accumulate later in evolution than the mediator mutations even though they produce similar fitness increases. This delay is consistent with the target for adaptive, cohesin-related mutations being smaller than the target for inactivating mediator. Genetic evidence suggests that the mutations in separase are mild loss-of-function mutations, but the effect of mutations in Smc1 and Smc3 are unclear. Mutations in these proteins can directly alter their interactions with kleisin, but any mutation that disrupts the essential biochemical activity of cohesin will be lethal. We argue that the number of mutations that change the regulation of the cohesin complex but not its essential activity is small, explaining the later accumulation of these mutations.

We argue that considering the target size for different mutations explains why we saw no mutations in Rec8. Since Rec8 and Scc1 have diverged substantially roughly a billion years, it may require multiple, simultaneous amino acid substitutions in Rec8 to improve its ability to hold mitotic sister chromosomes together. Even if single amino acid substitutions in Rec8 can improve its function in mitosis, there are unlikely to be many such mutations and the selective advantage conferred by individual mutations is likely to be modest. In contrast, the target size for inactivating mutations, such as those in mediator and the G1-to-S regulators, are large. If the mutations in *SMC1*, *SMC3*, and *ESP1* reduce some aspect of their function, the target size for mutations in these genes will be larger than the target size for mutations that improve Rec8’s mitotic function. Our results are consistent with other studies where loss-of-function mutations are the first step in adaptation in laboratory evolution experiments (Hottes et al., 2013; Koschwanez et al., 2013; Laan et al., 2015; Wildenberg and Murray, 2014).

Mutational target size is likely to explain why adaptive mutations often occur outside the gene whose product is being asked to perform a different function. When *E. coli* is experimentally evolved to use an enzyme that normally participates in proline synthesis, ProA, to catalyze a similar reaction in arginine synthesis, most of the adaptive mutations are in other genes of arginine synthesis pathway, not in ProA (Morgenthaler et al., 2019). Mutations outside the focal gene are also found when proteins are asked to perform the same function in a novel cellular environment. Thus *E. coli* adapts to use orthologs of the *folA* gene, which encodes dihydrofolate reductase, via mutations in genes responsible for protein degradation rather than mutations in the *folA* ortholog (Bershtein et al., 2015). We suggest that evolutionary changes in a protein’s function reflect a mixture of changes in its sequence and expression, changes in the proteins that it physically interacts with, and changes in other proteins that contribute to the biological function under selection. The number and diversity of these connections makes it difficult to predict the evolutionary trajectories that populations will follow as proteins are selected to perform new functions. Thus, evolutionary repair experiments are a strategy to learn more about the factors that regulate protein function and reveal previously unknown links between different functional modules.

Mutations in three functional modules, transcriptional mediator, chromosome cohesion, and cell cycle regulation improve the fitness and chromosome segregation of Rec8-expressing cells. We used double mutants to probe the interactions between these modules, scoring both fitness and sister cohesion. Most pairs of mutations interacted roughly additively for both phenotypes, with some exceptions: double mutations with *esp1-P15* were substantially fitter than the additive expectation and double mutations with *mbp1*Δ increased fitness but did not improve sister cohesion, suggesting that this mutation has effects on both sister cohesion and some other function. None of the adaptive mutations increase the total level of Rec8 in mitotically-arrested cells, suggesting that they likely alter the ability of Rec8-containing cohesion to form and maintain the linkages that hold sister chromosomes together.

Our work reveals a new regulatory link between sister cohesion and genome replication. The Rec8-expressing strain advances the timing of genome-wide origin firing and completes genome replication earlier than wild type. All the adaptive mutations we tested extend S phase and improve sister cohesion. We found deletion of the Cdk8 gene, separase mutation, and cohesin mutation all restore the pattern of origin firing towards that of wild type. This result is consistent with multiple populations acquiring mutations that inactivate genes (*MBP1*, *CLN2*, *SWI4*, and *SWI6*) that promote passage from G1 to S phase. We speculate that slowing genome replication can promote sister cohesion in the Rec8-expressing strain. We tested the causality of this linkage using mutants that reduce origin firing or slow replication forks: both manipulations improve the fitness and sister chromosome cohesion of Rec8-expressing cells, demonstrating that slower replication raises fitness. A recent study demonstrated that Mad2, the spindle checkpoint protein, regulates S phase by promoting translation of Clb5 and Clb6 (Gay et al., 2018), potentially explaining why *mad2*Δ also extends the S phase of Rec8-expressing cells. Our work demonstrates that mitotic sister chromosome cohesion can be improved by mutations that slow genome replication. The simplest explanation of this effect is that slower replication allows more time for Rec8-containing cohesin to associate with chromosomes either before or during the passage of the replication fork.

Our work leads to a number of questions. Why does expressing Rec8 advance the timing of origin firing while removing Scc1 has no effect? How do mutations in transcriptional mediator and cohesin-related genes affect replication? Are the effects of Rec8 on replication in the mitotic cycle related to its reported ability to stimulate genome replication in the meiotic cycle (Cha, 2000)? In early S phase, replication initiation is orchestrated by a series of molecular interactions: the proteins that activate DNA replication, like S-phase cyclin-dependent kinase (CDK) and Dbf4-dependent kinase (DDK), recruit several initiation factors to form an activated helicase complex (Bell and Labib, 2016). Cells can control the timing of origin firing by modifying the activity of these activators, limiting the dosage of replication initiation factors (Mantiero et al., 2011) or changing local chromatin structure which affects how easily these regulators can access a given origin (Aparicio, 2013; Boos and Ferreira, 2019). Further research is needed to determine whether Rec8 affects replication by increasing the expression of genes that control replication initiation or altering chromatin structure to make replication origins more accessible to initiation factors, or some combination of both. The combination of overexpressing four initiation factors and reducing chromatin compactness accelerates the firing of late replication origins, demonstrating that these factors can alter the dynamics of replication (Mantiero et al., 2011).

The most pressing question is how mutations in the transcriptional mediator complex, the major target of early adaptive mutations, alter the timing of genome replication and increase the fitness of Rec8-expressing cells. There are suggestions that transcriptional mediator is involved in genome replication. In budding yeast, genes of the Cdk8 complex have genetic interactions with genes that trigger replication initiation (*DBF4, DPB11, SLD3,* and *CDC7*) and core helicase components (*SLD5*) (Costanzo et al., 2016). In fission yeast (Banyai et al., 2017) and mammalian cells (Kohler et al., 2019), mutations of the Cdk8 complex are reported to alter genome replication, suggesting our finding that the Cdk8 module is involved in genome replication is not species specific, although the detailed mechanism remains unknown.

Overall, this evolution experiment shows that mutations outside the meiotic kleisin, Rec8, improve its ability to support mitotic chromosome segregation. We argue that the distinct functions of mitotic and meiotic kleisins evolved through a mixture of changes in kleisin itself and changes in other functional modules that regulate sister chromosome cohesion directly or indirectly. At least in laboratory experiments, the size and complexity of this molecular network provides a much larger target for mutations that alter the biological function of kleisin than the target presented by kleisin itself. We suggest that the functional divergence of paralogous proteins depends on a mixture of mutations in the paralogs and the proteins that they directly or indirectly interact with.

## Materials and Methods

### Yeast Strains, Plasmids, and Growth Conditions

All yeast strains are derivatives of W303, and their genotypes are listed in Supplementary File 2. For the yeast strain with a GFP labeled chromosome V and P*_GAL1_*-*SCC1*, yPH344 and yPH345 are haploid strains derived from a diploid strain that made from a cross between FY1456 (the same strain as K2789, a gift from Dana Branzei) and yPH36. yPH346 is a haploid strain derived from a diploid strain that made from a cross between FY1456 and yPH115. Strains carrying *REC8* integrated at the endogenous *SCC1* locus were generated by homologous recombination: A *REC8*-3xHA fragment was amplified from the plasmid pFA6a-*REC8*-3xHA-*KANMX4*, fused with 500 bp upstream and downstream DNA fragments of SCC1 coding sequence by PCR, and recombined with the *SCC1* genomic locus. Strains used in the evolution experiment were generated from yPH280 by plasmid shuffling (Lundblad and Zhou, 2001). Standard rich medium, YPD (1% Yeast-Extract, 2% Peptone, and 2% D-Glucose) was used for the evolution experiment. Growth conditions for each experiment are specified in the figure legends. Raffinose and Galactose were used at 2%. Benomyl was used at 30μg/ml. Cycloheximide was used at 35μg/ml. α-factor was used at 10μg/ml for *bar1* strains and at 100μg/ml for *BAR1* strains. Methionine was used at 8mM.

### Experimental Evolution

The haploid strain used in the evolution experiment was *MATα scc1*Δ pRS*414*-P*_SCC1_*-*REC8*-HA. To force yeast cells depend on Rec8 for mitotic growth, five clones of yHP280 (*MATα scc1*Δ pRS*414*-P*_SCC1_*-*REC8*-HA pRS416-*SCC1*) were cultured in YPD to lose pRS416-*SCC1* and cells without the *SCC1*-bearing plasmid were selected by the growth on 5-FOA plates. For each of the five clones, three independent 5-FOA resistant colonies were chosen, giving rise to the fifteen ancestral clones in the evolution experiment. Each ancestral clone was cultured in 3ml YPD to reach 10^8^ cells/ ml at 30°C and diluted 1:6000 into a tube with 3ml fresh YPD and incubated for 48 hrs (before generation 375) or 24 hrs (after generation 375). Each subsequent cycle used the same dilution. We estimated an effective population size of 6.3 × 10^5^ cells using this formula (Lenski et al.,1991), *N*_*e*_ = *N*_*o*_ × *g*, in which *N*_*o*_ is the initial population size (5 × 10^4^cells) and *g* is the number of generations, 12.6, during one cycle. After every ten cycles, 1ml culture was mixed with 500μl 80% Glycerol and frozen at −80°C. The evolution experiment was continued for 1750 generations.

### Fitness measurement by competition assay

An ancestral *P_SCC1_*-*REC8* strain that expressed a fluorescent protein, mCitrine, under the *ACT1* promoter was used as the reference strain (yPH447) in fitness competition assays with evolved populations and reconstructed strains carrying a single evolved mutation. For scoring fitness and sister cohesion in the same strain carrying a GFP-labeled Chr. V, a Rec8-expressing strain with P*_ACT1_*-*mChrerry* (yPH472) was used as the reference strain.

All sample strains and reference strain were grown to <10^7^cells/ml in YPD; cell density was measured using a Coulter Counter (Beckman Coulter). At the first time point, samples strains, either evolved strains or reconstructed strains carrying evolved mutations, were mixed with the reference strain at a ratio of 1:10. The initial cell mixture was diluted to 5×10^4^ cells/ ml in YPD and grown for 24 hours. At the second time point, the cell density was usually around 5×10^6^ cells/ml. Cultures were diluted to 5×10^4^ cells/ml to grow another 20-24 hours as the third time point. At each timepoint, 5×10^4^ cells of each mixed culture were transferred to single wells of a 96 well plate with U-shaped bottom for flow cytometry. A BD LSRFortessa FACS machine equipped with High Throughput Sampler was used to collect 30000 cells to quantify the ratio of the sample and the reference strain. The FACS data was analyzed using the FlowJo10.4.1 software. In addition to being mixed with sample strains, the reference strain was cultured separately to estimate the number of generations in an experiment. Each experiment was conducted in technical triplicates, and the fitness of each sample strain was measured in three independent experiments.

To calculate relative fitness, *w*, of each sample strain to the reference strain, we followed this formula: *w* = 1 + *s*,

*s* is selection coefficient: 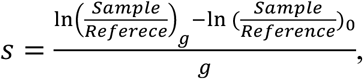

in which *g* is the number of generations and 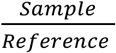 is the ratio between a sample strain and a reference strain (Desai et al., 2007).

### Chromatin Immunoprecipitation and qPCR

We followed the protocol of calibrated chromatin immunoprecipitation (Makrantoni et al., 2019) to precipitate chromosome-bound kleisins. First, cell quantities were measured by multiplying culture volumes by optical density at 600 nm (OD_600_) and this product is referred to as O.D. units. 20 O.D. of *S. cerevisiae* cells were crosslinked with 1% formaldehyde for 30minutes at 25°C. Each *S. cerevisiae* cell pellet was mixed with 15 O.D. of crosslinked *Schizosaccharomyces pombe* cells that expressed an epitope tagged version of the Scc1 homolog (*RAD21*-HA). The inclusion of the fission yeast cells served as control to normalize technical variations between samples. This mixture was resuspended in ChIP lysis buffer A (50mM HEPES-KOH at pH7.5, 0.1M NaCl, 1mM EDTA, 150mM NaCl, 1% TritonX-100, 0.1% Sodium Deoxycholate, 1x protease inhibitor (Roche) and further lysed by bead beating (BioSpec Products) with 0.5mm glass beads (Biospec Products). To shear chromatin, a Covaris S220 instrument was used with the following program: peak incident power:175, duty factor: 10%, cycle per burst: 200, treatment time: 250. After shearing, the cell lysate was centrifuged at 16000g at 4°C for 20 minutes to collect the supernatant containing protein-bound, sheared chromatin. To pull down the fraction of chromatin bound by kleisin, 15 μl pre-washed dynabeads ProteinG (Invitrogen) and 7.5μl anti-HA antibody (12CA5, Invitrogen) were added in 1ml lysate and incubated at 4°C overnight. After immunoprecipitation, the beads were washed with ChIP wash buffer I (50mM HEPES-KOH at pH7.5, 0.1M NaCl, 1mM EDTA, 274mM NaCl, 1% TritonX-100, 0.1% Sodium Deoxycholate), ChIP wash buffer II (50mM HEPES-KOH at pH7.5, 0.1M NaCl, 1mM EDTA, 500mM NaCl, 1% TritonX-100, 0.1% Sodium Deoxycholate), ChIP wash buffer III (10mM Tris/HCl pH8.0, 0.25M LiCl, 1mM EDTA, 0.5% NP40, 0.5% Sodium Deoxycholate), and TE (10mM Tris/HCl pH8.0, 1mM EDTA).

To process the sample for quantitative PCR, immunoprecipitated chromatin and a 1:100 dilution of the input chromatin were separately recovered by boiling with a 10% Chelex-100 resin (BioRad) before treating with 25 μg/ml Proteinase K at 55 °C for 30 minutes. Samples were boiled again to inactivate Proteinase K, centrifuged and the supernatant was subjected to qPCR on ABI 7900 using PerfeCTa SYBR Green FastMix ROX (Quanta BioSciences). The sequences of primers used for qPCR are listed in Table S3. To calculate the enrichment of pull-down DNA in total input chromatin,

We used the following formula: 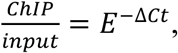

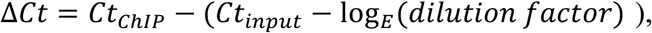 in which *E* is primer efficiency and *dilution factor* is 100. Enriched % of input is 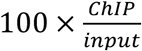. At each cohesin binding site, the final enriched % of input is calibrated to the enriched % of input at a peri-centromeric site of the *S. pombe* genome.

For ChIP-Seq, chromatin was immunoprecipitated as described above, and purified chromatin was subjected to DNA end repair and dA tailing to make sequencing library as described (Makrantoni et al., 2019). Samples were sequenced on a MiniSeq with 75 base paired-end reads (Illumina, San Diego, CA). Scripts and workflows used to create ChIP-Seq are stored on the github repository (https://github.com/PhoebeHsieh-yuying).

### Cell cycle progression by flowcytometry

Yeast strains with *P_GAL1_*-*SCC1* were grown in YEP containing 2% Galactose to log phase. To synchronize populations in G1, cells were washed and diluted in YEP containing 2% Raffinose and 100 μg/ml alpha-factor for 2 hours. G1 synchronization was confirmed by checking that the percentage of cells that had formed mating projections (shmoos) was over 90% using a light microscope (MICROPHOT-SA, Nikon). To restart the cell cycle without *SCC1* transcription, cells were washed with YEP containing 50 μg/ml Pronase (Sigma-Aldrich) twice and resuspended in YPD containing 50 μg/ml Pronase at 30°C. 1ml of cells were collected and fixed by 70% Ethanol at G1 and several timepoints after growth in YPD according to the figure legend of each experiment. Subsequently, fixed samples were treated with 0.4mg/ml RNase A (Sigma-Aldrich) at 37°C overnight, followed by 1mg/ml Proteinase K (Sigma-Aldrich) treatment at 50°C for 1 hour. DNA was stained with 1μM Sytox Green solution (Invitrogen). Prior to flow cytometry, the stained cells were sonicated for 30 seconds with 70% intensity by BRANSON Ultrasonics Sonifier S-250. A total of 10,000 cells were collected using a BD LSRFortessa FACS machine (Becton Dickinson). The FACS data was analyzed using FlowJo10.4.1. To quantify the fraction of replicating cells in the population at 30 minutes after release from the G1 block, the Watson (Pragmatic) model built in the cell cycle analysis tools of FlowJo was used to calculate the portion of cells that had completed replication (G2 peak), were undergoing genome replication (S phase), and were still in G1.

### Sister chromosome cohesion assay and Microscopy

Strains with *tetO_112_* array integrated at the *URA3* locus and *tetR*-*GFP* were used for assaying cohesion between sister chromatids (Uhlmann and Nasmyth, 1998). To examine sister chromosome cohesion in one cell cycle, sample preparation followed the same procedure as the cell cycle progression experiment excepts that cells were released from their G1 arrest into YPD containing 30μg/ml benomyl to arrest them once they reached mitosis. At each time point, cells were fixed in 4% paraformaldehyde for 15 minutes, washed, and stored in a storage solution (1.2M Sorbitol, 0.1M KH_2_PO_4_/K_2_HPO_4_) at 4°C. Images were taken with a 100x objective on a Nikon inverted Ti-E microscope with a Yokagawa spinning disc unit and an EM-CCD camera (Hamamatsu ImagEM); GFP was excited with a 488 nm laser with 25% laser power. For each image, a z-stack was taken with 41 z-steps spaced 0.5 μm apart. Images were analyzed by the Fiji distribution of ImageJ (Schindelin et al., 2012). For each experiment, at least 100 cells are analyzed. Three independent experiments were conducted for each strain.

### Measurement of protein abundance and protein stability

The protein-containing extracts from cell pellets were prepared by NaOH lysis (Kushnirov, 2000) and analyzed on Western Blots. Protein extracts were resuspended with SDS sample buffer (10mM Tris pH6.8, 2% SDS, 10% glycerol, 0.004% bromophenol blue, and 2% *β*-mercaptoethanol) and boiled at 100°C for 5mins. Rec8 and Scc1 were all tagged with 3xHA at the C-terminus of coding sequences and anti-HA antibody (3F10, Roche) was used to detect the abundance of kleisin proteins. The abundance of Hxk1 was monitored as a loading control using an anti-Hxk1 antibody (USBiological Life Sciences, H2035-01). SuperSignal West Dura reagent (Thermo Scientific) was used for developing chemiluminescent signal. Chemiluminescent signals were detected by an Azure Sapphire Biomolecular Imager and quantification of protein abundance was analyzed by ImageStudioLite.

To measure protein stability in mitosis, cells were arrested at metaphase in YPD containing 30μg/ml Benomyl for 2 hours and treated with cycloheximide at 35 μg/ml to inhibit protein synthesis. 1ml cells were collected for to prepare samples for Western Blotting at the indicated timepoints after adding cycloheximide.

### Whole Genome Sequencing and Analysis

Genomic DNA was prepared as described (Koschwanez et al., 2013). DNA sequencing libraries were prepared using the Illumina Nextera DNA library Prep kit as described (Baym et al., 2015). The sequencing was done on an Illumina HiSeq 2500 with 125 base paired-end reads or Illumina NovaSeq with 150 base paired-end reads. Whole genome sequencing data was processed as described (Koschwanez et al., 2013). The Burrow-Wheeler Aligner (bio-bwa.sourceforge.net) was used to map DNA sequences to the *S. cerevisiae* reference genome r64, downloaded from *Saccharomyces* Genome Database (www.yeastgenome.org). The resulting SAM (Sequence Alignment/Map) file was converted to a BAM file, an indexed pileup format file, using the samtools software package (samtools.sourceforge.net). GATK (www.broadinstitue.org/gatk) was used to realign local indels, and Varscan (varscan.sourceforge.net) was used to call variants. Mutations were identified using an in-house pipeline (github.com/koschwanez/mutantanalysis) written in Python. Variants that differ between the ancestral and evolved genome >10%, a threshold above average sequencing error, are called as mutations, and any mutation present in >90% of sequencing reads in the evolved genome is defined as fixed mutation. In our pipeline, mutations can be found in both coding and non-coding sequences. In this study, we focused on mutations that cause non-synonymous substitution in the coding sequence. Scripts and workflows used to find evolved mutations are stored on the github repository (https://github.com/PhoebeHsieh-yuying).

### Generation of Reconstructed Strains

Individual mutations from evolved strain were engineered into the targeted locus of the ancestral strain by homologous recombination. A DNA fragment containing targeted gene with the desired mutation, a selection maker (*HpHMX4* or *HIS3MX4*), and 300 bp downstream of the targeted gene was made by PCR. To measure fitness effect of an evolved mutation in the ancestor, this DNA fragment was transformed into yPH280. The presence of the desired mutation was confirmed by Sanger sequencing. To measure fitness effect of reconstructed mutations in the Rec8-expressing background, yeast cells were cultured in YPD to lose pRS416-*SCC1* and cells without *SCC1* plasmid were selected by the growth on 5-FOA plates.

To measure the effect of these reconstructed mutations on cell cycle progression and sister cohesion, the same transformation procedure was done in the *P_GAL1_*-*SCC1 P_SCC1_*-*REC8* strain with GFP-labeled chromosome V and phenotypes were assayed in the presence of glucose.

### Whole Genome Replication Profile

Yeast cells were arrested in G1 with 2μg/ml α-factor and low pH as described (Rosebrock, 2017) and follow the same procedure allowing cells enter cell cycle as the flowcytometry experiments. 1ml cells were collected separately for DNA content analysis and genomic DNA extraction at the following time points: 0, 10, 20, 30, 40, 50, 60 minutes after G1 arrest. Whole genome sequencing libraries were made as previously described. Sequencing was done on an Illumina NovaSeq with 150 base paired-end reads. Two separate experiments were done for each strain.

The analysis of genome replication profile was done as described (Saayman et al., 2018); (Fumasoni and Murray, 2019). Reads mapping and CNVs detection were processed as (Koschwanez et al., 2013). We followed the script in (Fumasoni and Murray, 2019) to analyze change in the CNVs at multiple timepoints during S phase to generate whole genome replication profile. First, read-depth of every 100bp window is normalized to the medium read-depth of a sequenced genome to control for sequencing variation between samples. To allow intra-strain comparison at multiple timepoints, the normalized read-depth was further scaled to the medium of DNA content obtained by flow cytometry to generate relative coverage to the corresponding G1 genome. The resulting coverage was then averaged across multiple 100bp windows and a polynomial data smoothing filter (Savitsky-Golay) was applied to the individual coverage profiles to filter out noise. Replication timing T_rep_ is defined as the time at which 50% of the cells in the population replicated a given region of the genome, which is equivalent to an overall relative coverage of 1.5x, since 1x corresponds to an unreplicated region and 2x to a fully replicated one. The replication timing T_rep_ was calculated by using linear interpolation between the two time points with coverage lower and higher than 1.5x to compute the time corresponding to 1.5x coverage. Final T_rep_ were then plotted relative to their window genomic coordinates. Scripts and workflows used to generate whole genome replication profiles are stored on the github repository (https://github.com/marcofumasoni).

## Supplementary Figures

**Figure S1.**
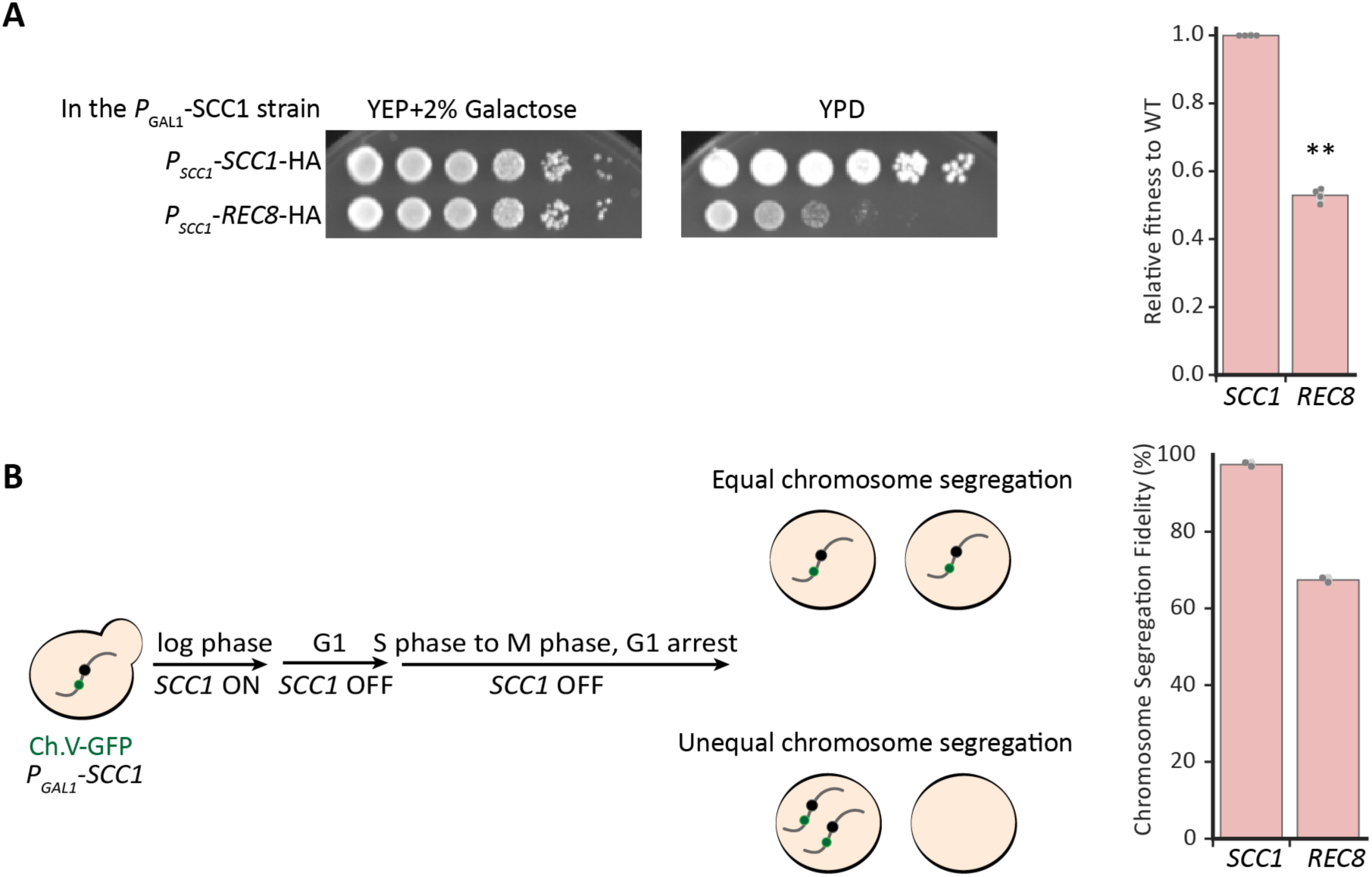
Mitotic growth and chromosome segregation of the *P_GAL1_-SCC1 P_SCC1_-REC8* strain. We used a *P_GAL1_-SCC1 P_SCC1_-REC8* strain to examine the effects of acutely expressing Rec8 as the sole kleisin. Cells were propagated in galactose-containing medium, arrested in G1, and then released into glucose-containing medium to repress Scc1. **(A)** The *P_SCC1_*-*REC8* strain grows poorly when *SCC1* expression is turned off. Left: Cells were grown in YEP containing 2% galactose to the same density and serially diluted on YEP containing 2% galactose or 2% glucose, in which the *GAL1* promoter was repressed by glucose. Right: Relative fitness of a *P_GAL1_-SCC1 P_SCC1_-REC8* strain to that of wild type in YPD. The darker gray points represent the values of three biological replicates and the thinner gray bar represents one standard deviation on each side of the mean of these measurements. (two-tailed Student *t* test, ** *p* < 0.01) **(B)** The fidelity of chromosome segregation of the Rec8-expressing strain is 30% lower than that of wild type. *P_GAL1_-SCC1 P_SCC1_-REC8* cells were grown in YEP containing 2% Galactose to log phase, transferred to YEP containing 2% raffinose and alpha-factor to repress *SCC1* expression and arrest them in G1, prior to release into YPD to resume cell cycle with *SCC1* expression repressed. Once cells had entered S phase, alpha-factor was added again to prevent cells entering a second cell cycle. Chromosome segregation fidelity was measured as the fraction of G1-arrested cells in a population showing one GFP dot, representing one copy of chromosome V, after one mitotic cell division. At least 100 cells were imaged in each experiment. The darker gray points represent the values of two biological replicates and the thinner gray bar represents one standard deviation on each side of the mean of these measurements.

**Figure S2.**
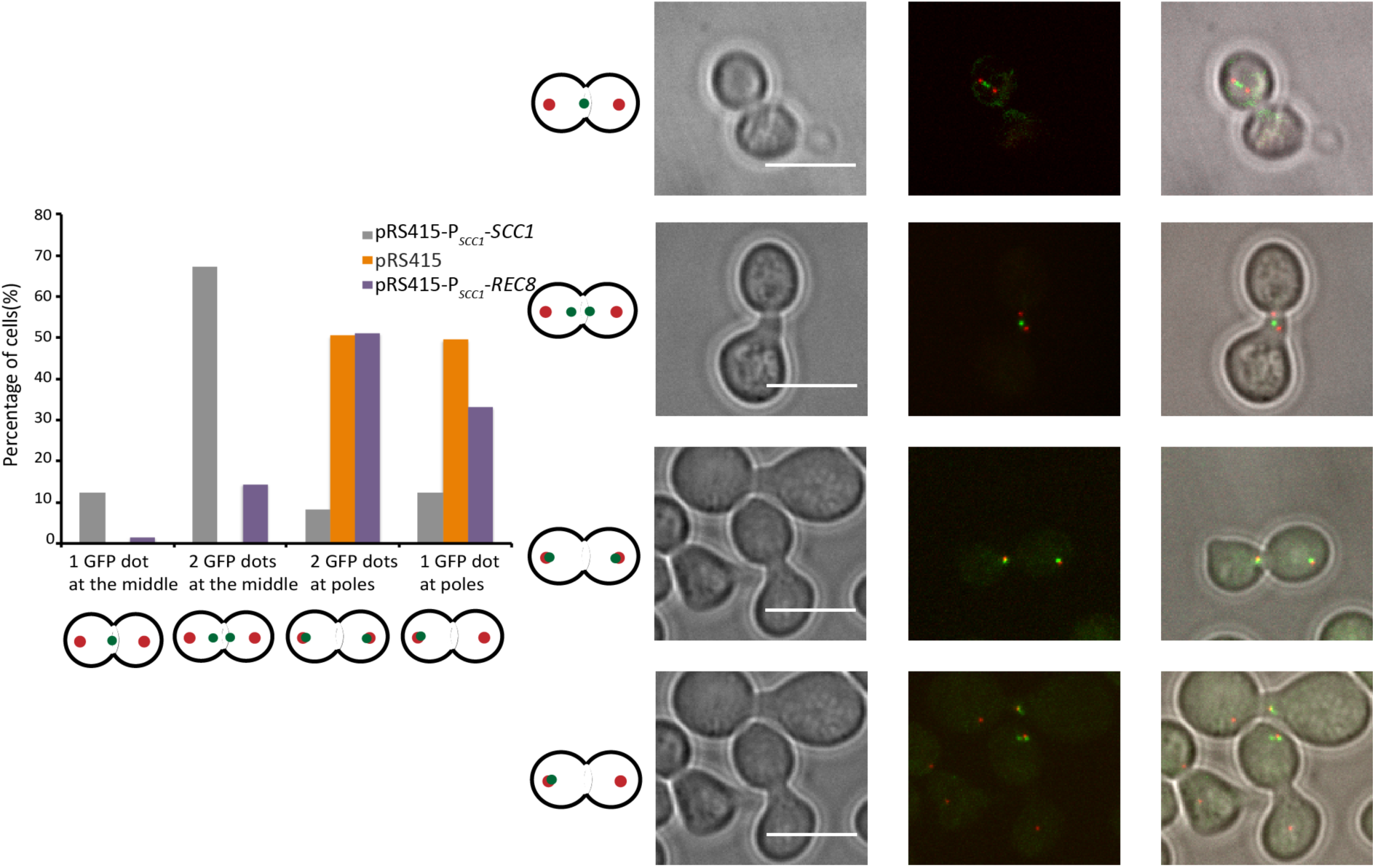
Sister kinetochore biorientation is perturbed in *P_SCC1_-REC8* cells. The yeast strain P*_MET_*-*CDC20-3xHA* P*_GAL1_*-*SCC1-3xHA CEN15*::*LacO* P*_CUP1_*-*GFP-LacI SPC42*-*mCherry* was transformed with a pRS415-based plasmid of *P_SCC1_*-*SCC1*, *P_SCC1_*-*REC8*, or an empty plasmid. Cells were cultured in CSM-Met-Leu containing galactose to log phase and switched to CSM-Met-Leu containing raffinose and alpha-factor to be synchronized in G1. Then, to repress the *SCC1* expression and arrest cells in metaphase, cells were released into YEP containing glucose and methionine for one cell cycle. The centromere of chromosome XV was marked by GFP and spindle pole bodies were labeled by *SPC42*-*mCherry*. Cells showing one or two GFP dots in the middle of two spindle pole bodies represent bi-oriented sister kinetochores under tension exerted by the spindle. The lack of sister chromosome cohesion leads to sister chromosomes of chromosome IV separating in prometaphase, resulting either in one GFP dot at each spindle pole body or two GFP dots at one of the spindle pole bodies. At least 100 cells were imaged and analyzed in each population. Illustrative microscopy images are shown in the right; the scale bar is 10μm.

**Figure S3.**
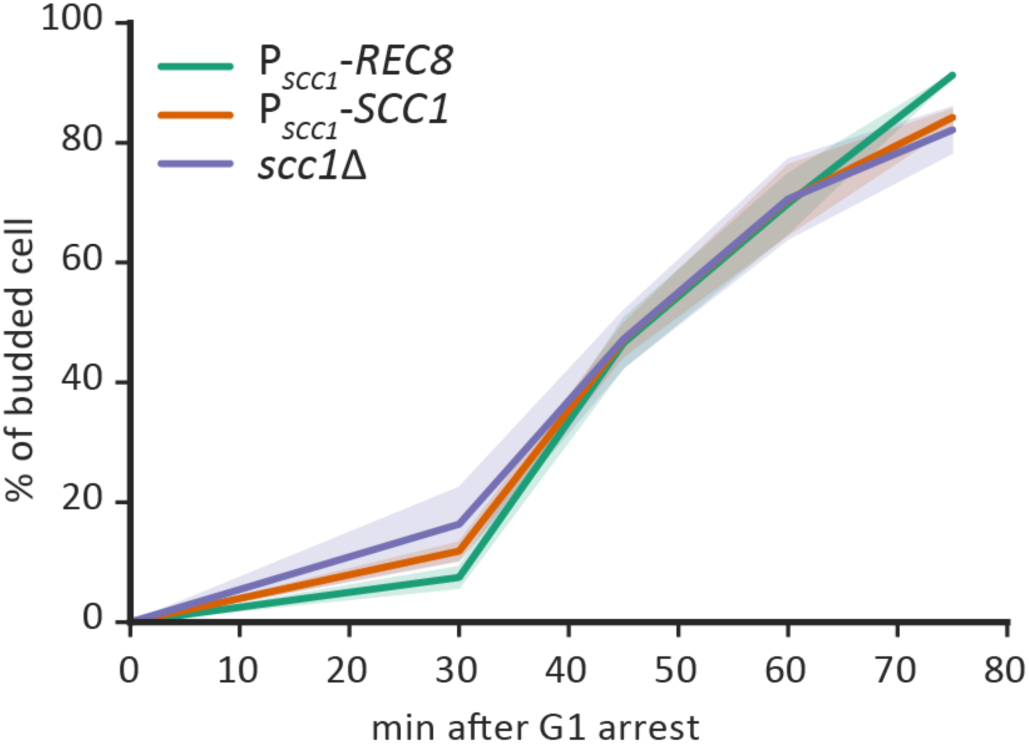
The budding index of the Rec8-expressing strain is similar to that of wild type and the *scc1*Δ strains. Cells were arrested in G1 and then released as described in Fig. 1C. The y-axis shows the fraction of budded cells in a population, measured as budding index. At least 100 cells were examined at each time point for each experiment. The mean (solid line) and standard deviation (shaded region) of three biological replicates for each population are shown.

**Figure S4.**
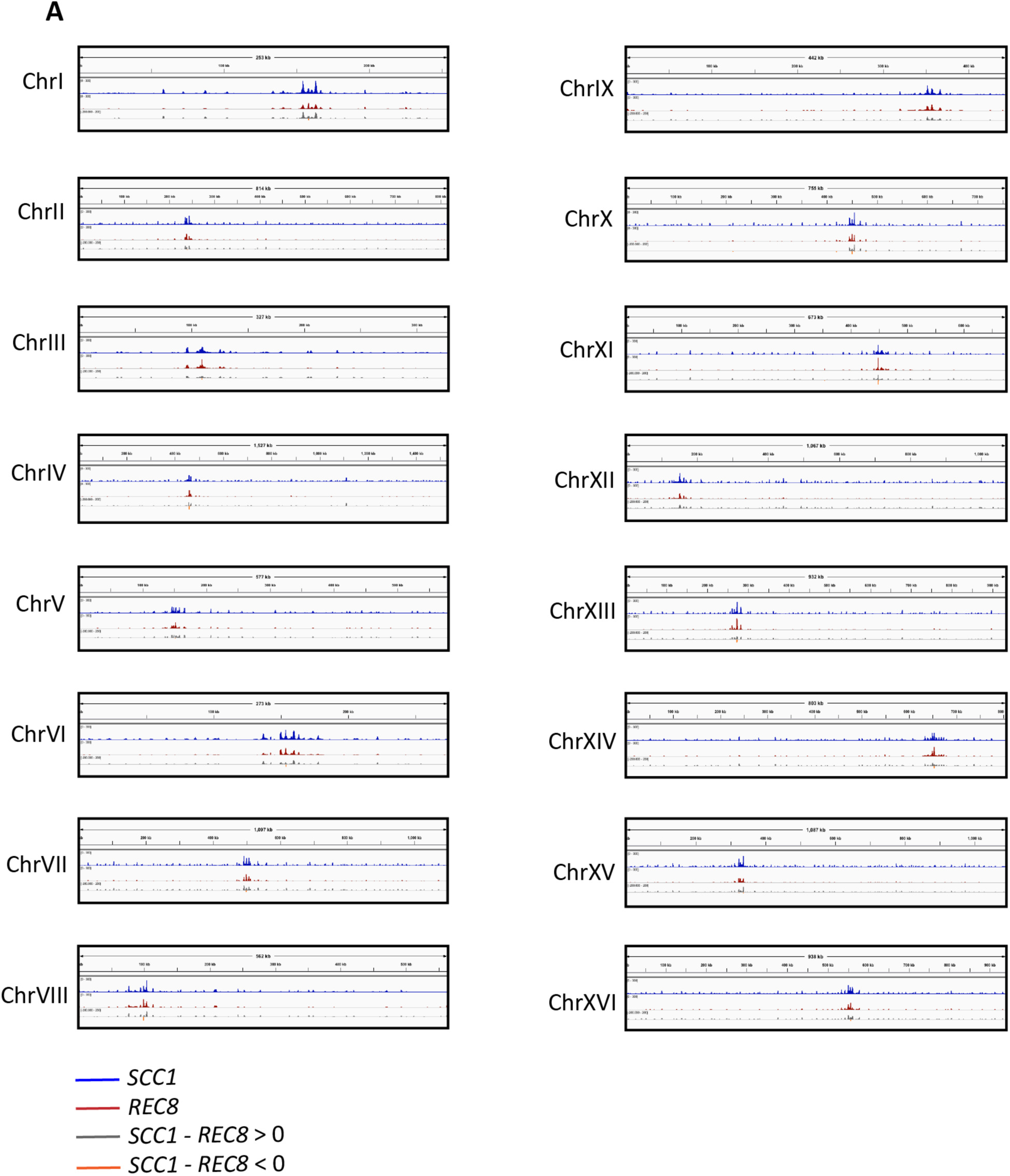
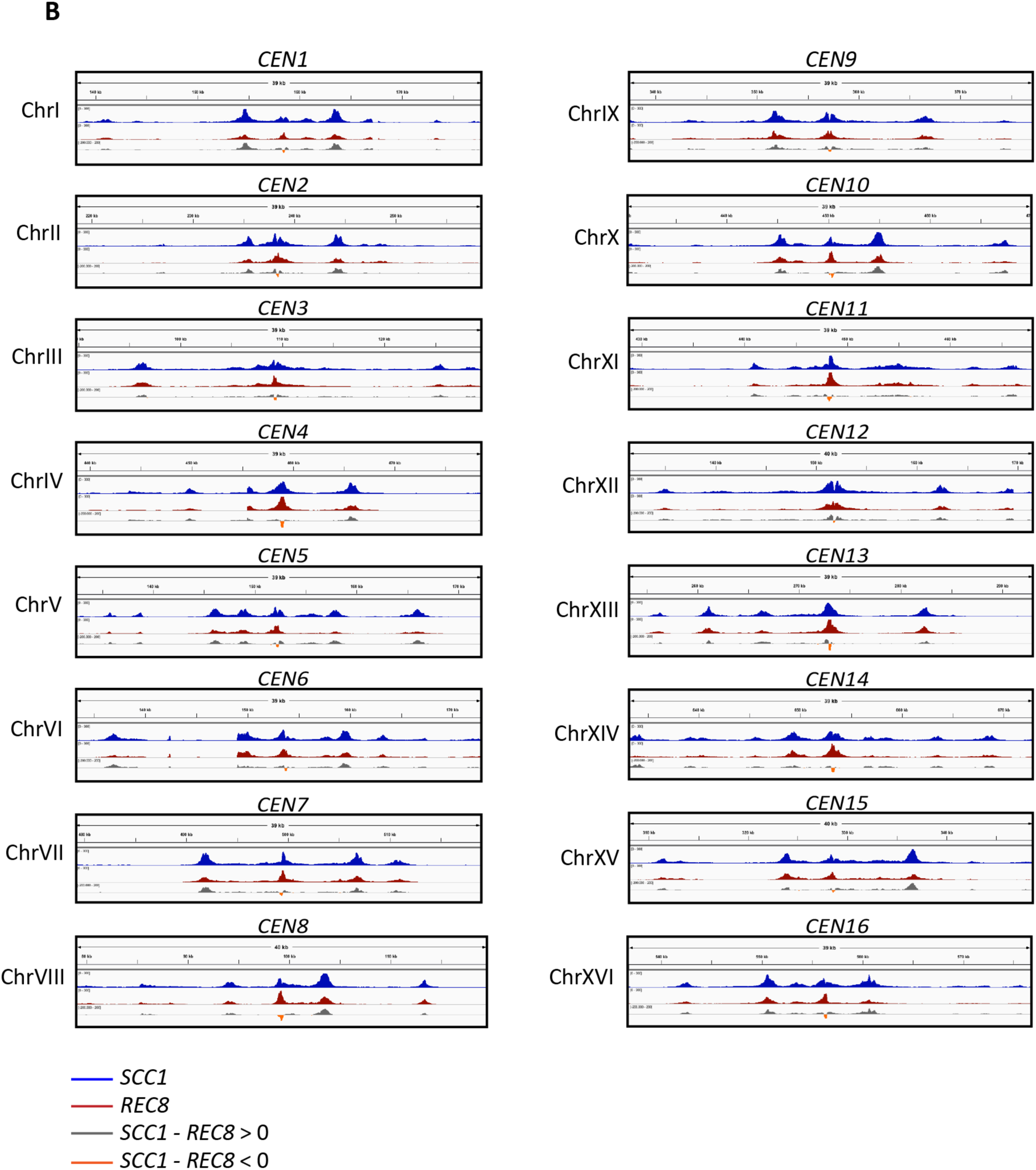
The genome wide enrichment of Rec8 is lower than that of Scc1. The ChIP-Seq data for all 16 chromosomes (chromosomes III and XV are also shown in Fig. 2D). Read depths calibrated to an internal control of the *S. pombe* genome are shown on the y-axis as reads per million (RPM, 0-300). The enrichment of Scc1and Rec8 is shown in blue and red respectively. The difference in the read depth between Scc1 and Rec8 is shown in the last track of each panel, in gray where Scc1’s signal is higher than Rec8’s and in orange where Rec8’s signal is higher than Scc1’s. (A) ChIP-Seq data of individual chromosome. (B) ChIP-Seq data of individual centromeres extending 20 kb on either side of the centromere. Images were prepared using the Integrated Genome Viewer (Robinson et al., 2011).

**Figure S5.**
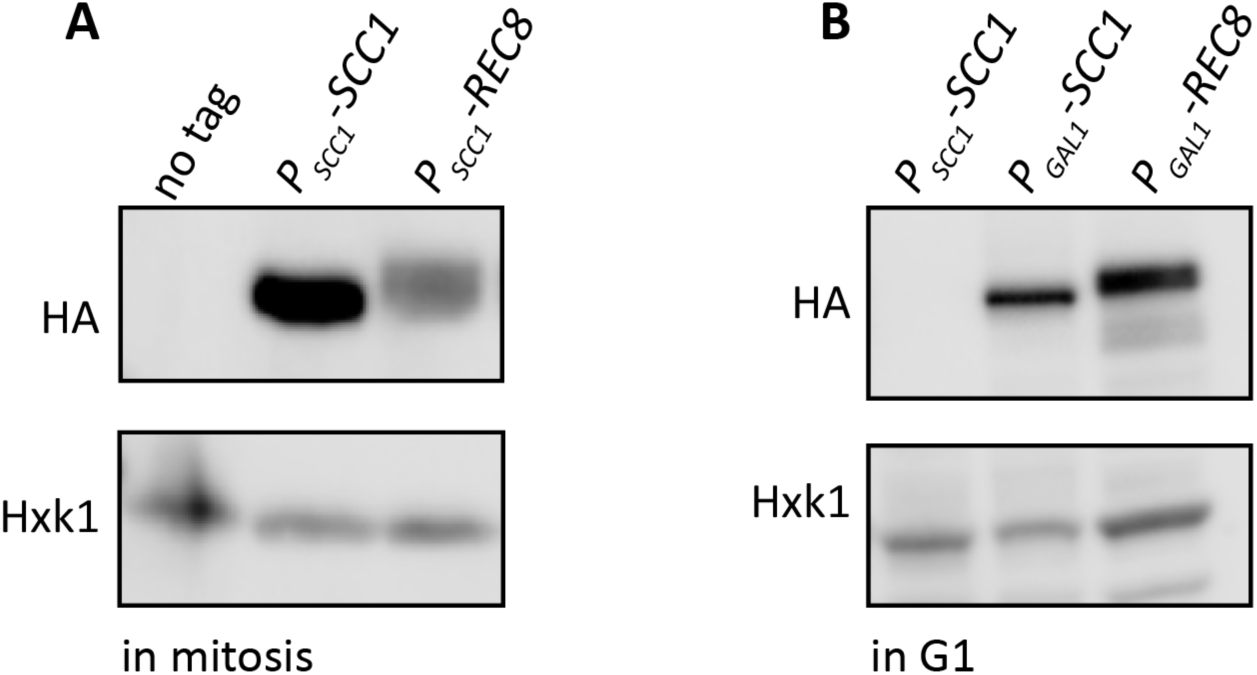
Protein levels of Scc1 and Rec8 in cell extracts processed for ChIP experiments. (A) Protein levels of two kleisins in mitosis. Cells were processed as described in Fig. 2C and cell extracts were obtained by alkaline lysis prior to analysis by Western Blotting. Kleisin proteins were detected by anti-HA antibody and hexokinase (Hxk1) was used as a loading control. (B) Protein levels of two ectopically-expressed kleisins in G1. Cells were processed as described in Fig. 2E and cell extracts were obtained by alkaline lysis prior to analysis by Western Blotting. The *P_SCC1_*-*SCC1*-HA strain was used as a negative control because the endogenous *SCC1* gene is not expressed in G1. Hexokinase (Hxk1) was used as a loading control.

**Figure S6.**
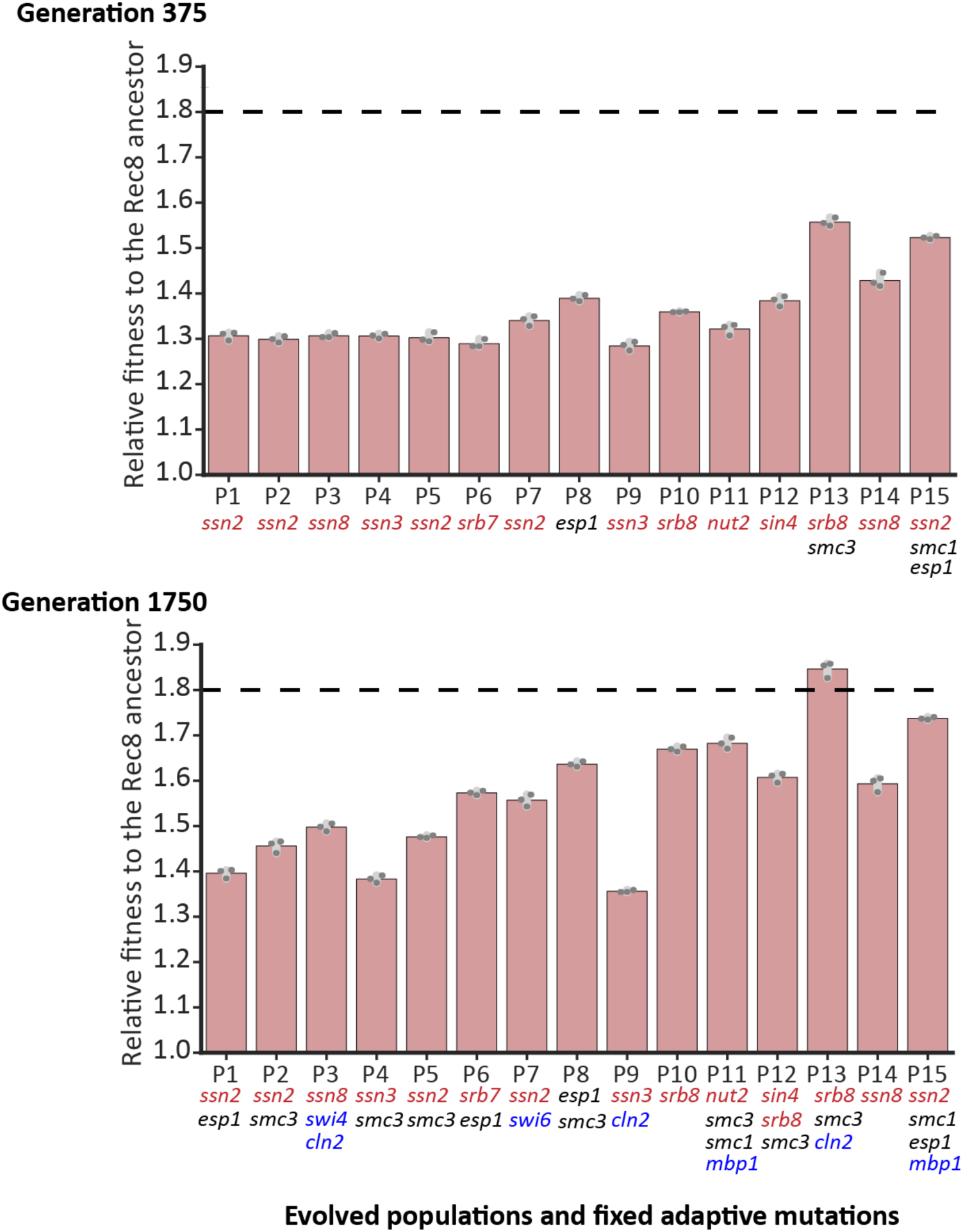
The fitness of fifteen evolved populations relative to the Rec8-expressing ancestor at generation 375 and generation 1750. The fitness of fifteen evolved populations at generation 375 (upper panel) and at generation 1750 (lower panel) is shown. The darker gray points represent the values of three biological replicates and the thinner gray bar represents one standard deviation on each side of the mean of these measurements. Genes in the three functional modules that mutate in at least six evolved populations at generation 1750 are shown. Mutations in the different functional modules are color-coded: transcriptional mediator complex (red), G1-to-S cell cycle regulators (blue), and cohesin and its regulators (black). The black dashed line shows the fitness of wild type relative to the Rec8-expressing ancestor.

**Figure S7.**
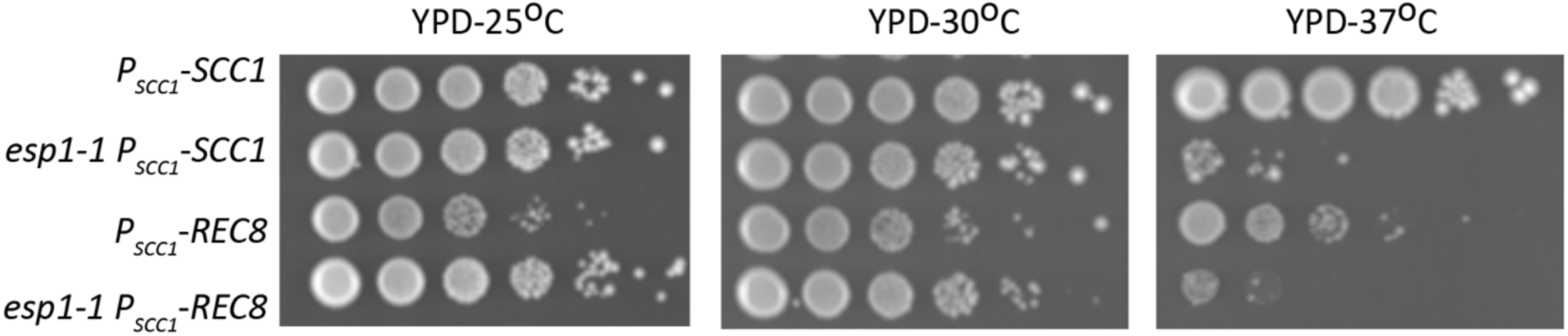
*esp1-1* improves the growth of the Rec8-expressing strain at 30°C. *esp1-1*, the known temperature-sensitive mutation that inactivates separase activity (Ho et al., 2015) was introduced to the Rec8-expressing strain by sporulating a heterozygous diploid strain (*P_SCC1_*-*SCC1*/*P_SCC1_*-*REC8 ESP1*/*esp1-1*). Haploid progeny carrying four different genotypes (*SCC1*, *SCC1 esp1-1*, *REC8*, and *REC8 esp1-1*) were selected. These four strains were subjected to serial dilutions and spotted on YPD. Their growth was measured at 25°C, 30°C, and 37°C.

**Figure S8.**
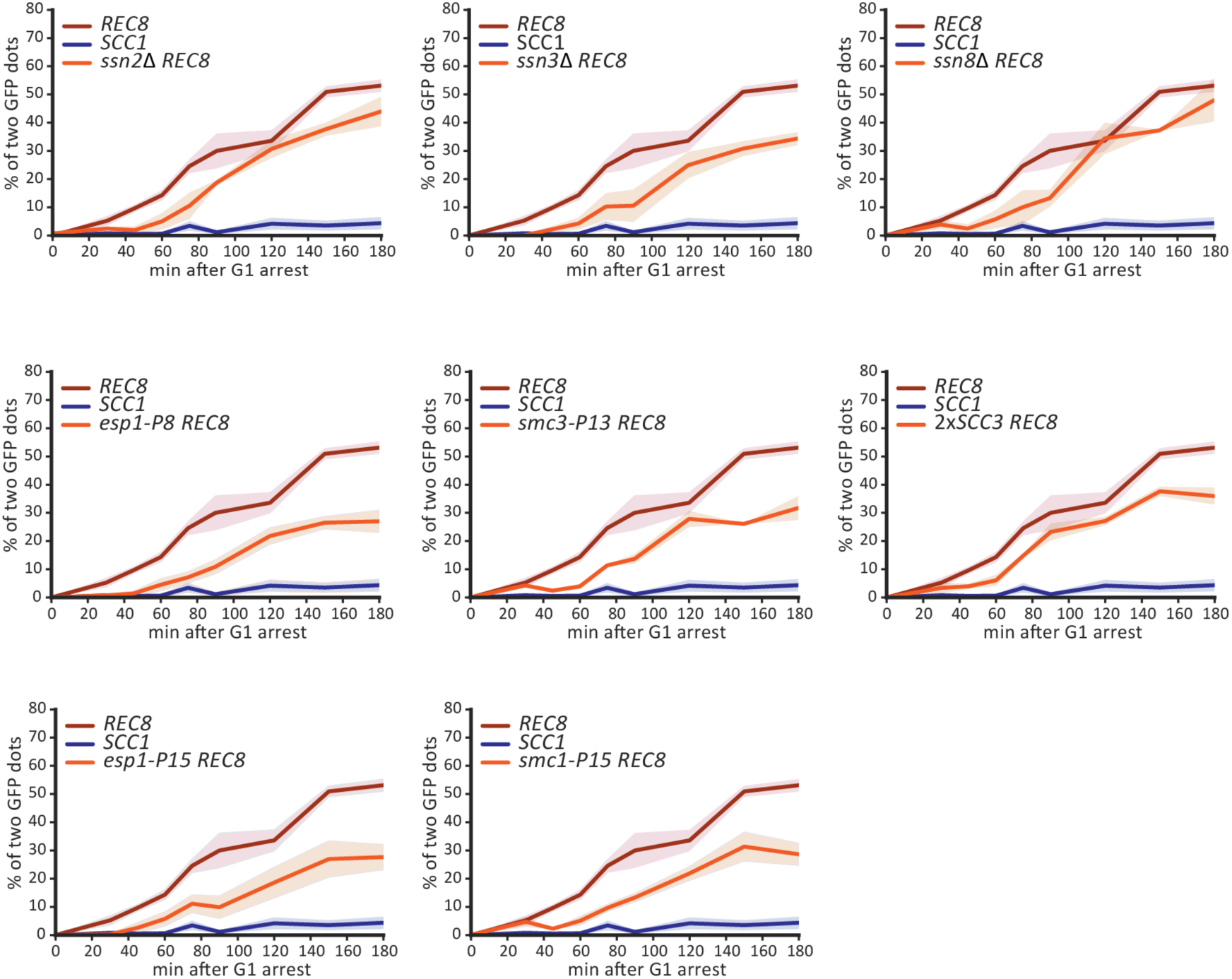
Individual reconstructed mutations partially improve sister cohesion. The time courses of sister chromosome separation for the experiment shown in Fig. 5A, which presents the data at 150 minutes after release from the G1 arrest. At least 100 cells were imaged at each time point for each experiment. The mean and standard deviation of three biological replicates are shown in the solid line and shading, respectively.

**Figure S9.**
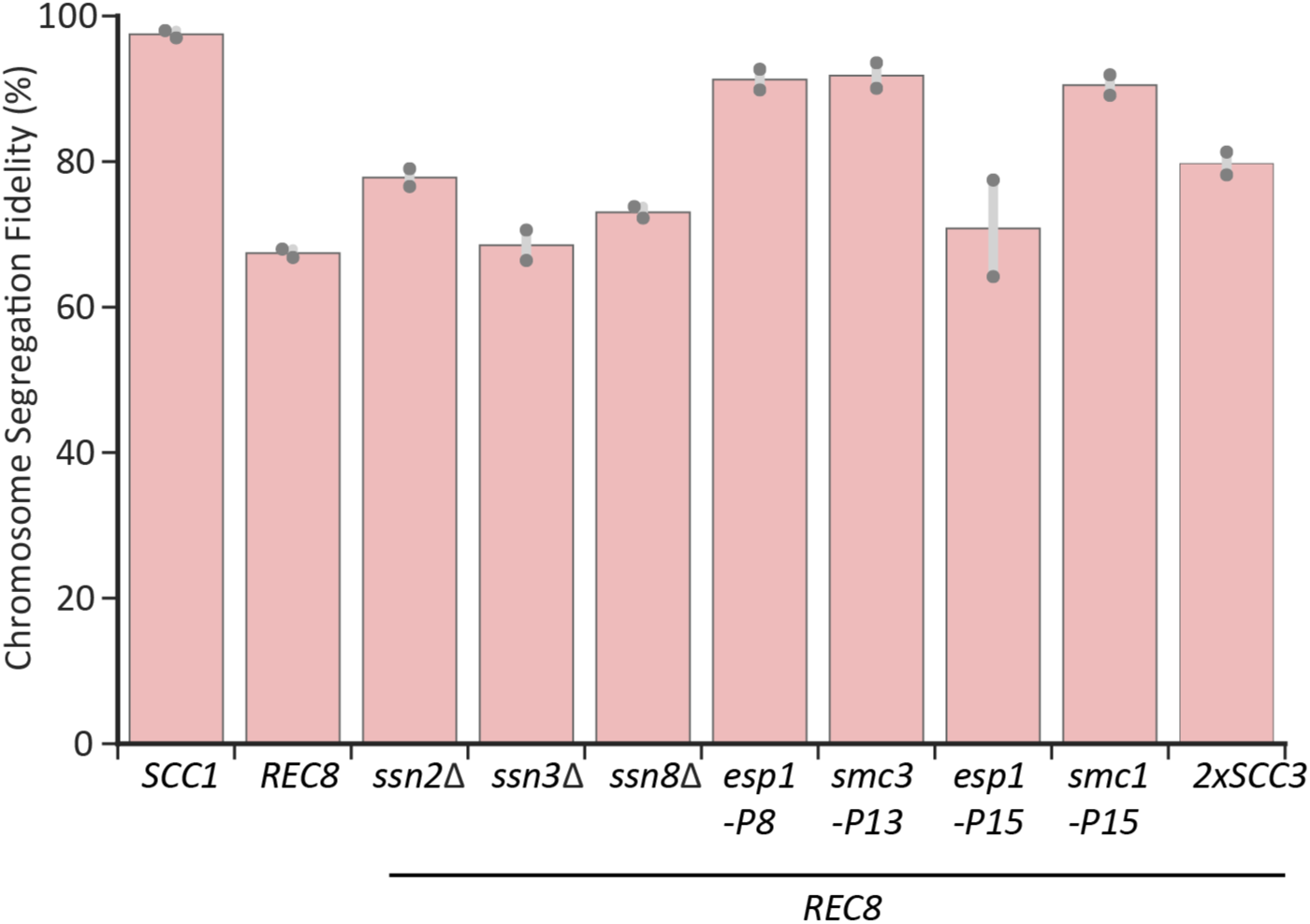
Individual reconstructed mutations increase chromosome segregation fidelity of the Rec8-expressing strain. Cells were prepared as in Fig. S1B to examine the fidelity of chromosome segregation in a single mitotic cell division. Gene deletions for three components of the Cdk8 complex were used to approximate the effect of the mutations of these genes found in evolved populations. At least 100 cells were imaged in each experiment. The darker gray points represent the values of two biological replicates and the thinner gray bar represents one standard deviation on each side of the mean of these measurements.

**Figure S10.**
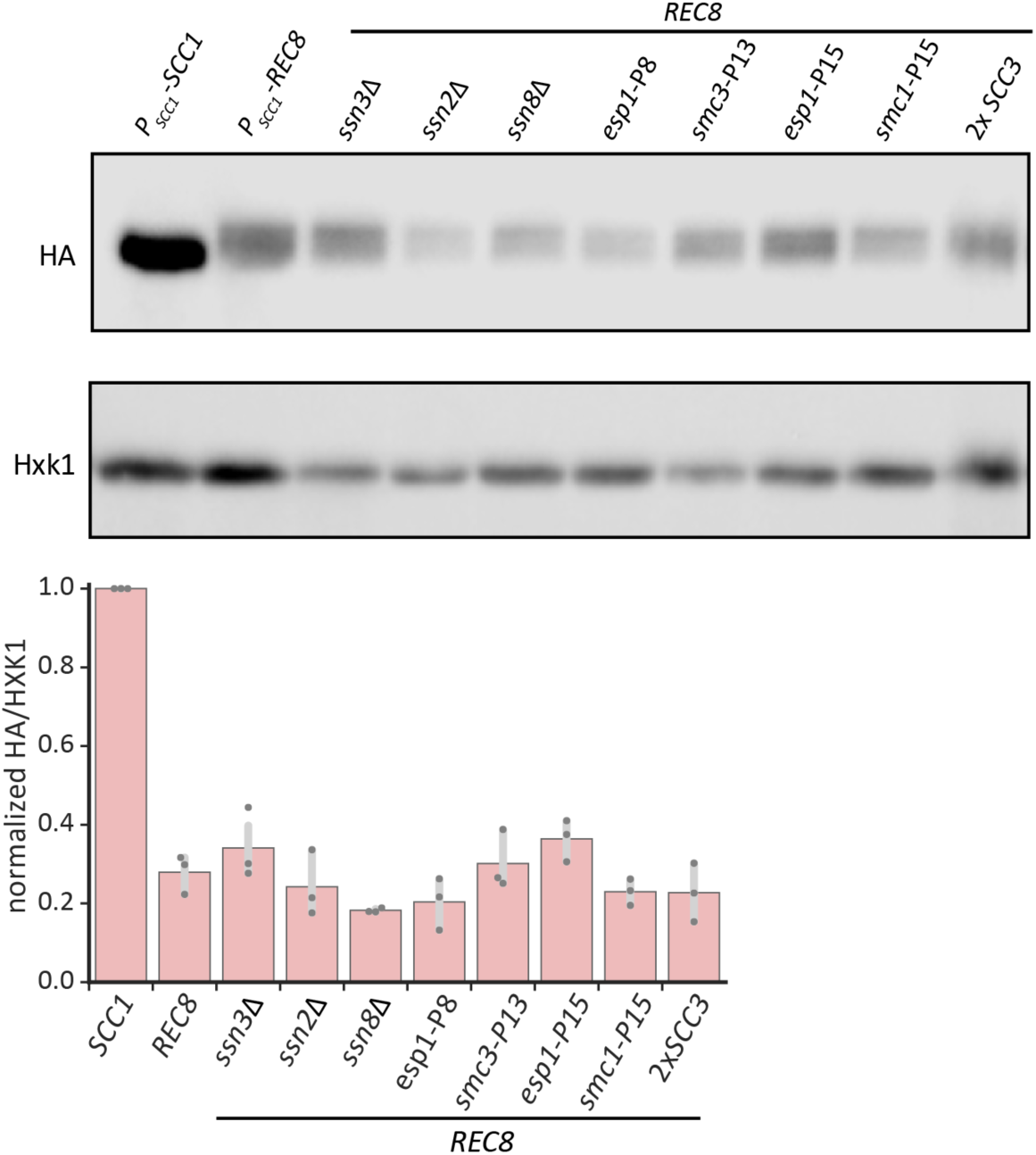
Individual reconstructed mutations do not alter the Rec8 protein level in mitosis. Strains with individual reconstructed mutations were synchronized in G1, released into cell cycle, and then arrested in mitosis in YPD containing benomyl. Protein samples were collected by alkaline lysis and analyzed by Western Blotting. Both Scc1 and Rec8 were tagged with 3xHA at their C termini and anti-HA antibody was used for their detection. Hxk1 (hexokinase) was used as loading control. In the bar graph, the darker gray points represent the values of three biological replicates and the thinner gray bar represents one standard deviation on each side of the mean of these measurements.

**Figure S11.**
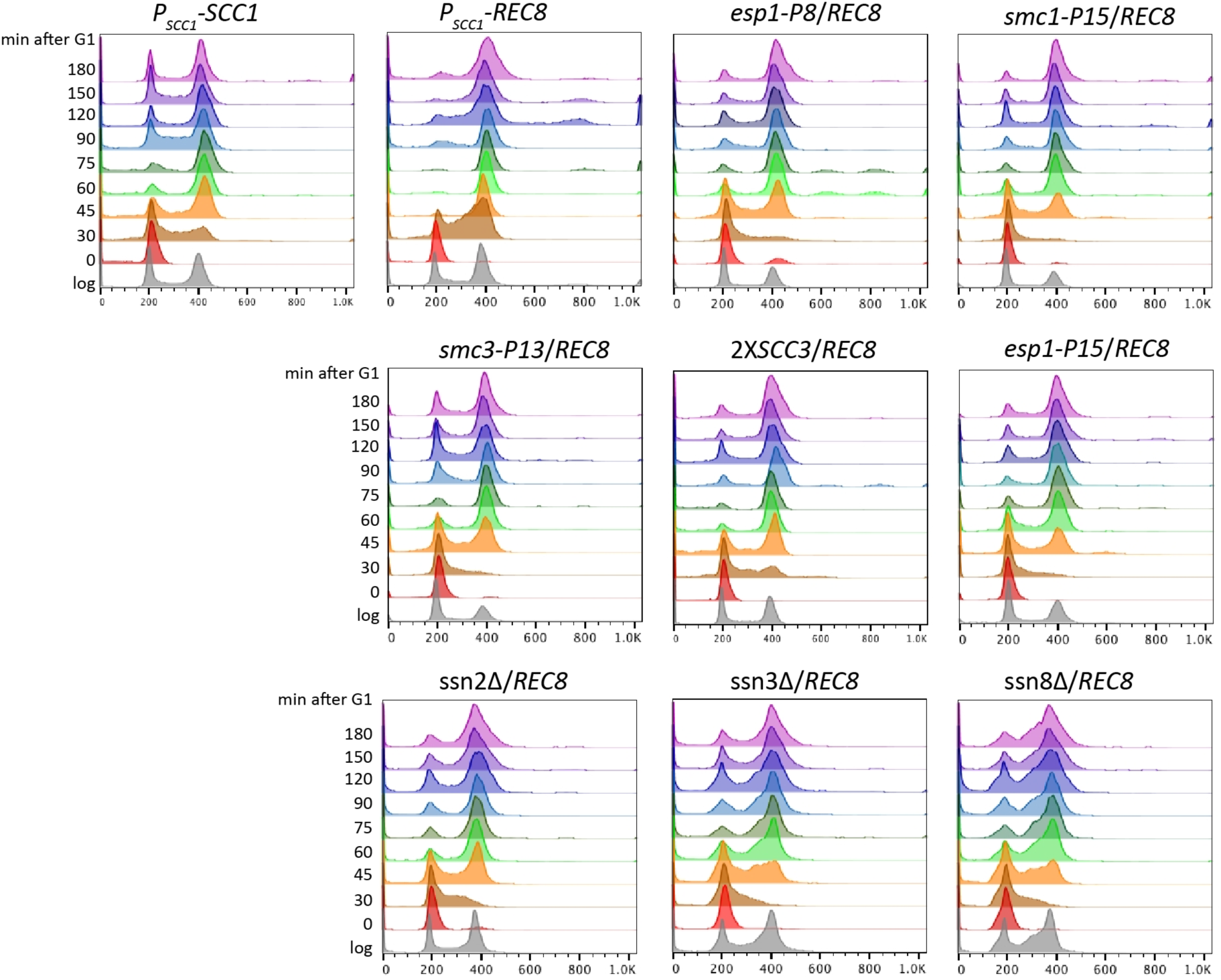
Individual reconstructed mutations partially restore the cycle progression profile of the Rec8-expressing strain towards that of wild type. The full time courses for the experiments summarized in Fig. 6A, which shows the time points from 0 to 75 minutes after release from the G1 arrest. Individual cohesin-related mutations, deletion of genes encoding the Cdk8 complex, and two integrated copies of *SCC3* were engineered separately into the *P_SCC1_*-*REC8 P_GAL1_*-*SCC1* background. Cells were allowed to proceed through a synchronous cell cycle as in Fig. 1D and were collected for fixation every 15 or 30 minutes following release from a G1 arrest to analyze their DNA content.

**Figure S12.**
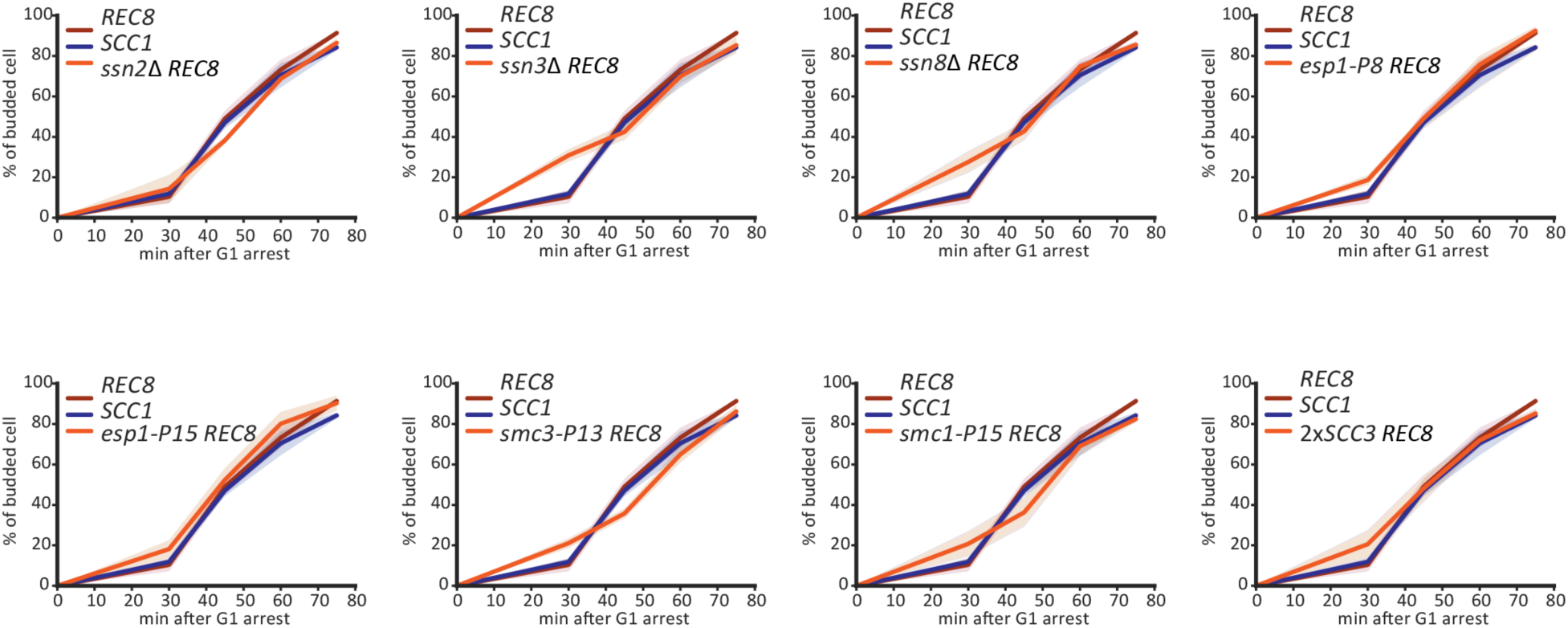
Individual reconstructed mutations don’t delay the onset of cell cycle in Rec8-expressing cells. Cells were arrested in G1 and then released as described in Fig. 6A. The y-axis shows the fraction of budded cells in a population, measured as the budding index. Strains carrying single reconstructed mutation (*ssn2*Δ, *ssn3*Δ, *ssn8*Δ, *esp1-P8*, *esp1-P15*, *smc3-P13*, *smc1-P15*, and two copies of *SCC3*) are compared with wild type and the Rec8-expressing strain. The mean (solid line) and standard deviation (shaded region) of three biological replicates for each strain are shown.

**Figure S13.**
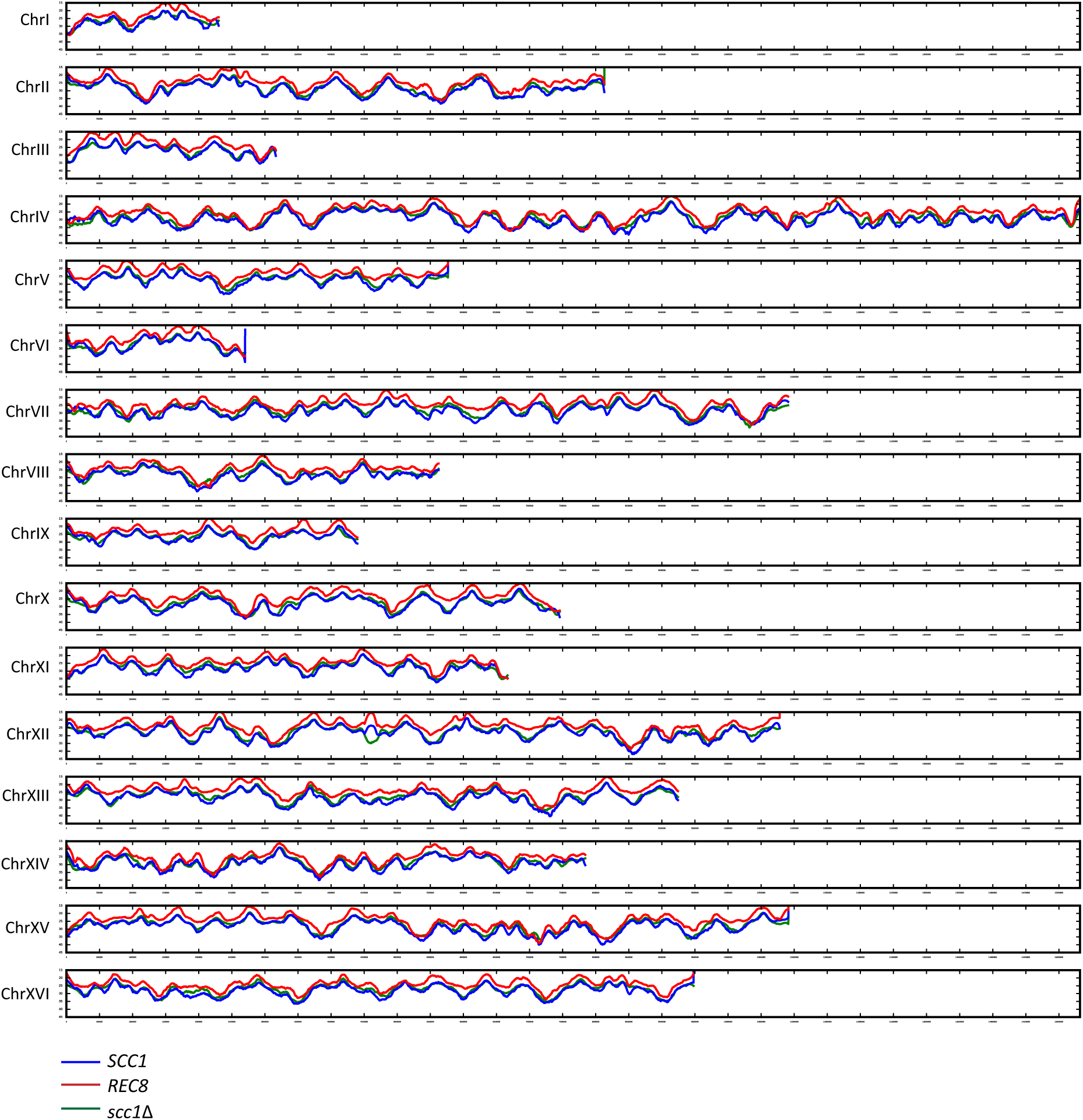
The whole genome replication profiles of wild type, the Rec8-expressing strain, and the *scc1*Δ strain. The mean replication profile of two experiments is shown. The replication profile of each strain is color-coded (*SCC1* in blue, *REC8* in red, and *scc1*Δ in green) and arranged by the order of chromosome. The y-axis represents T_rep_, the time at which 50% of cells in a population completes replication at a given genomic locus (See Materials and Methods for detailed analysis).

**Figure S14.**
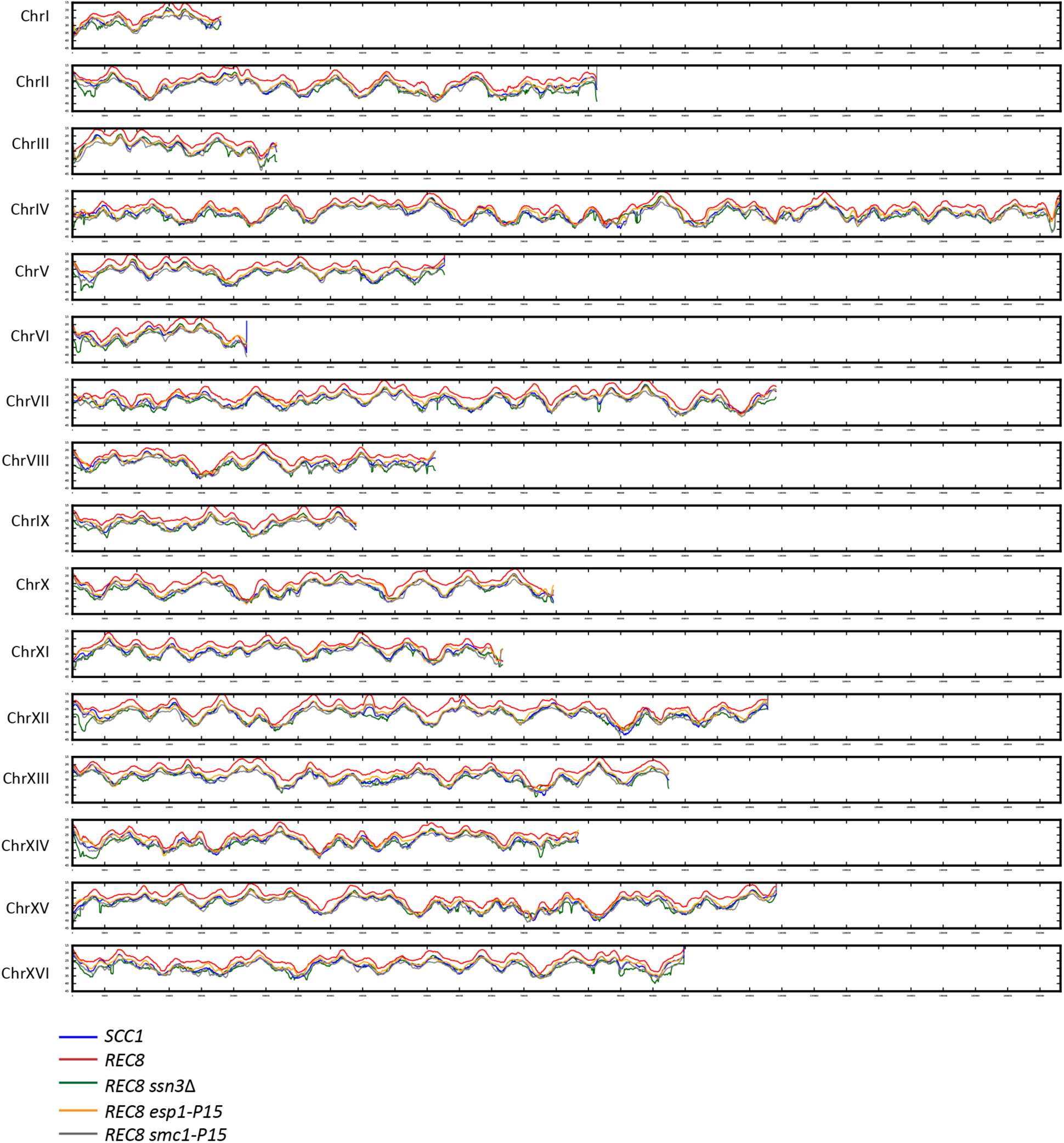
The whole genome replication profiles of the Rec8-expressing strain and reconstructed strains carrying a single evolved mutation, *ssn3*Δ, *esp1-P15*, or *smc1-P15*. These replication profiles are the data shown in Fig. 6E. The mean replication profile from two experiments is color-coded by strain and arranged by order of chromosome. The y-axis represents T_rep_, the time at which 50% of cells in a population completes replication at a given genomic locus (See Materials and Methods for detailed analysis).

**Figure S15.**
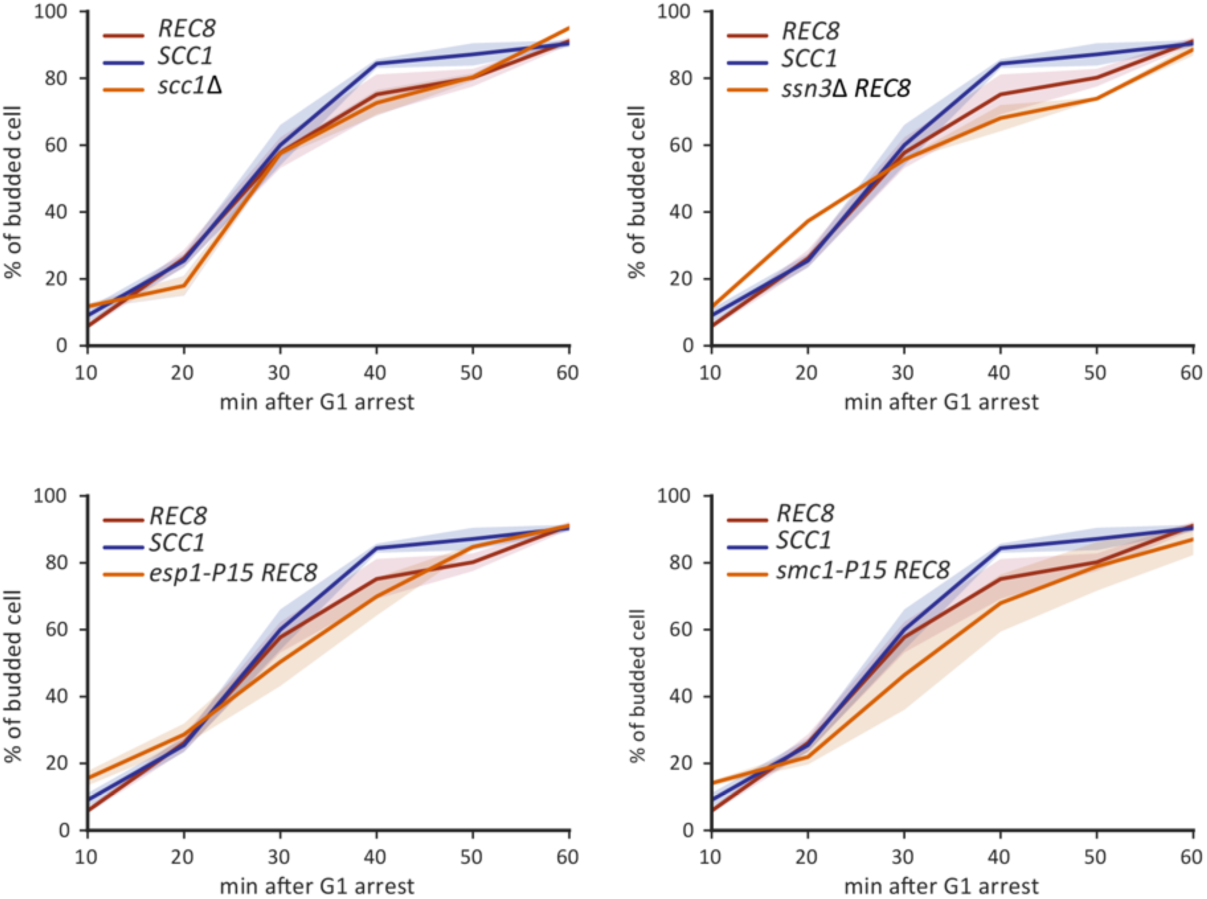
The budding index of yeast strains that are processed for replication profiling. Cells were arrested in G1 and then released as described in Fig. 6E, S13, and S14. The y-axis shows the fraction of budded cells in a population, measured as budding index. The mean (solid line) and standard deviation (shaded region) of two biological replicates for each strain are shown.

**Figure S16.**
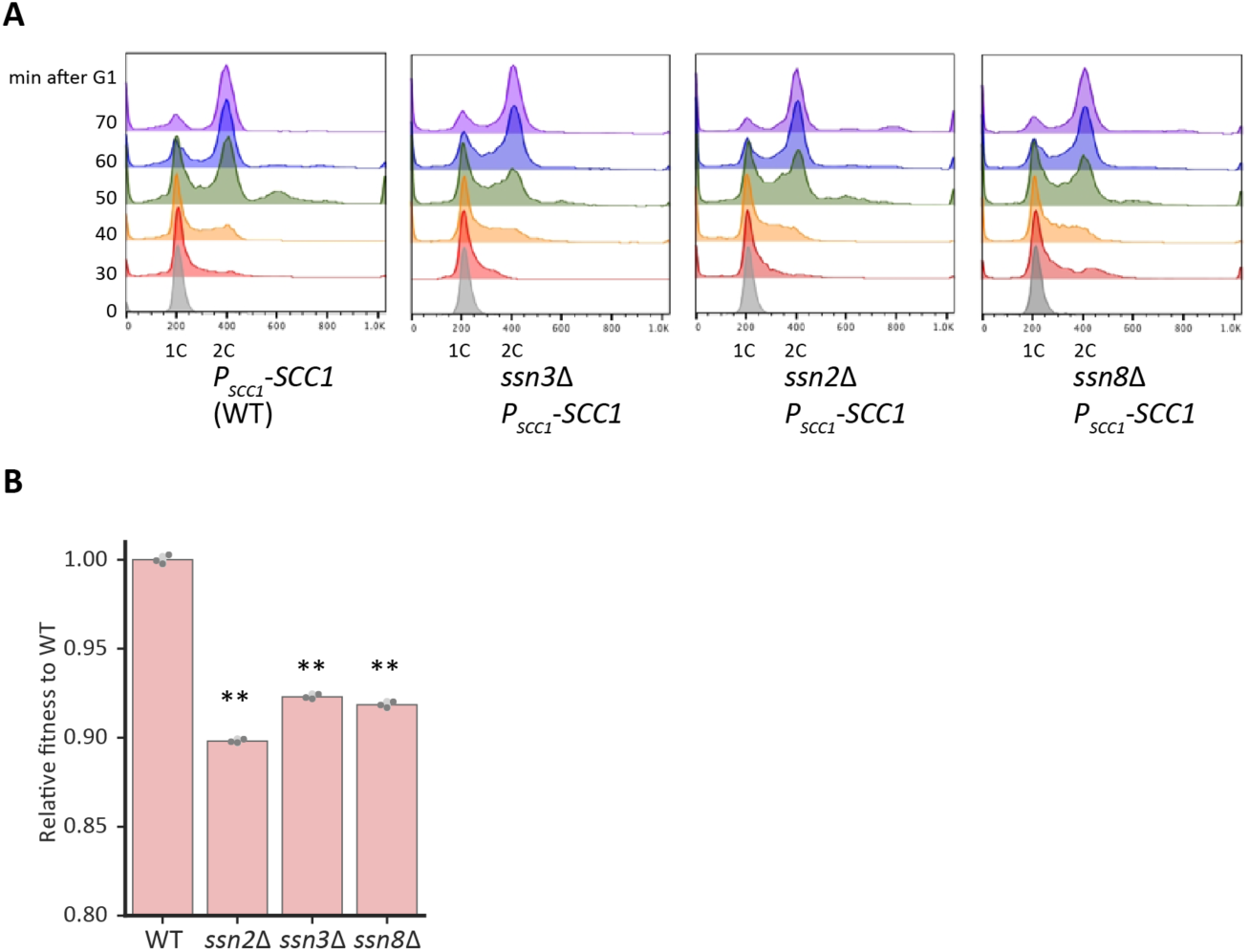
Deletion of genes encoding the Cdk8 complex slightly slows genome replication and cause 8-11% cost in wild type. **(A)** The cell cycle progression profiles of *ssn2*Δ, *ssn3*Δ, or *ssn8*Δ strains compared to a wild type control after release from a G1 arrest. (**B)** The fitness of *ssn2*Δ, *ssn3*Δ, or *ssn8*Δ strains relative to wild type, measured by competitive fitness assay. The darker gray points represent the values of three biological replicates and the thinner gray bar represents one standard deviation on each side of the mean of these measurements. The statistical significance between data from wild type and each mutant strain was calculated by two-tailed Student *t* test, ** *p* < 0.01.

**Figure S17.**
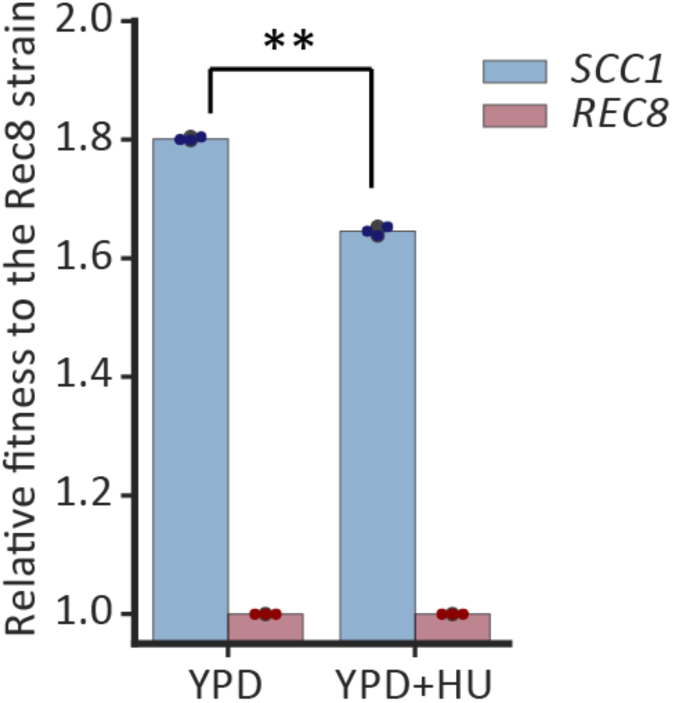
Hydroxyurea decreases the fitness difference between the Rec8-expressing strain and wild type. The fitness of wild type strains relative to the Rec8-expressing strain were measured in YPD and YPD containing 12.5mM hydroxyurea (HU). The colored points represent the values of three biological replicates and the darker gray bar represents one standard deviation on each side of the mean of these measurements. (two-tailed Student *t* test, ** *p* < 0.01)

## Acknowledgements

We thank Dana Branzei for providing yeast strain, Nichole Wespe and Marco Fumasoni for help with analysis of whole genome sequencing data, and Bauer Core Facility at Harvard for help with experiments. We thank Angelika Amon, Steve Bell, Jun-Yi Leu, Thomas LaBar, Andrea Giometto, Marco Fumasoni, and Hung-Ji Tsai for critical feedback on the manuscript. We thank members of the Murray lab and Marston lab for useful discussion. This work was supported by the following grants: Wellcome Senior Fellowship 107827, Welcome Centre core grant 203149, and NIH RO1-GM43987 and the NSF-Simons Center for the Mathematical and Statistical Analysis of Biology (#1764269 (NSF) & #594596 (Simons)).

## Author contributions

YPH, conception and design, acquisition of data, analysis and interpretation of data, drafting and revising the manuscript. VM, acquisition of ChIP-Seq, manuscript discussion and revision. DR, analysis of ChIP-Seq. ALM, manuscript discussion and revision. AWM, conception and design, interpretation of data, drafting and revising the manuscript.

## Reference

Aakre, C.D., Herrou, J., Phung, T.N., Perchuk, B.S., Crosson, S., and Laub, M.T. (2015). Evolving new protein-protein interaction specificity through promiscuous intermediates. Cell 163, 594–606.

Allen, B.L., and Taatjes, D.J. (2015). The Mediator complex: a central integrator of transcription. Nat Rev Mol Cell Biol 16, 155–166.

Amoutzias, G.D., He, Y., Gordon, J., Mossialos, D., Oliver, S.G., and Van de Peer, Y. (2010). Posttranslational regulation impacts the fate of duplicated genes. Proc Natl Acad Sci U S A 107, 2967–2971.

Aparicio, O.M. (2013). Location, location, location: it’s all in the timing for replication origins. Genes Dev 27, 117–128.

Azvolinsky, A., Dunaway, S., Torres, J.Z., Bessler, J.B., and Zakian, V.A. (2006). The S. cerevisiae Rrm3p DNA helicase moves with the replication fork and affects replication of all yeast chromosomes. Genes Dev 20, 3104–3116.

Banyai, G., Szilagyi, Z., Baraznenok, V., Khorosjutina, O., and Gustafsson, C.M. (2017). Cyclin C influences the timing of mitosis in fission yeast. Mol Biol Cell 28, 1738–1744.

Baym, M., Kryazhimskiy, S., Lieberman, T.D., Chung, H., Desai, M.M., and Kishony, R. (2015). Inexpensive multiplexed library preparation for megabase-sized genomes. PLoS One 10, e0128036.

Bell, S.P., and Labib, K. (2016). Chromosome Duplication in Saccharomyces cerevisiae. Genetics 203, 1027–1067.

Bershtein, S., Serohijos, A.W., Bhattacharyya, S., Manhart, M., Choi, J.M., Mu, W., Zhou, J., and Shakhnovich, E.I. (2015). Protein Homeostasis Imposes a Barrier on Functional Integration of Horizontally Transferred Genes in Bacteria. PLoS Genet 11, e1005612.

Boos, D., and Ferreira, P. (2019). Origin Firing Regulations to Control Genome Replication Timing. Genes (Basel) 10.

Brar, G.A., Hochwagen, A., Ee, L.S., and Amon, A. (2009). The multiple roles of cohesin in meiotic chromosome morphogenesis and pairing. Mol Biol Cell 20, 1030–1047.

Buonomo, S.B., Clyne, R.K., Fuchs, J., Loidl, J., Uhlmann, F., and Nasmyth, K. (2000). Disjunction of homologous chromosomes in meiosis I depends on proteolytic cleavage of the meiotic cohesin Rec8 by separin. Cell 103, 387–398.

Cha, R.S.W., B.M.; Keeney, S.; Dekker, J.; Kleckner, N. (2000). Progression of meiotic DNA replication is modulated by interchromosomal interaction proteins, negatively by Spo11p and positively by Rec8p. Genes Dev 14, 493.

Chothia, C., Gough, J., Vogel, C., and Teichmann, S.A. (2003). Evolution of the protein repertoire. Science 300, 1701–1703.

Cobbe, N., and Heck, M.M. (2004). The evolution of SMC proteins: phylogenetic analysis and structural implications. Mol Biol Evol 21, 332–347.

Costanzo, M., VanderSluis, B., Koch, E.N., Baryshnikova, A., Pons, C., Tan, G., Wang, W., Usaj, M., Hanchard, J., Lee, S.D., et al. (2016). A global genetic interaction network maps a wiring diagram of cellular function. Science 353.

Desai, M.M., Fisher, D.S., and Murray, A.W. (2007). The speed of evolution and maintenance of variation in asexual populations. Curr Biol 17, 385–394.

Donaldson, A.D., Raghuraman, M.K., Friedman, K.L., Cross, F.R., Brewer, B.J., and Fangman, W.L. (1998). CLB5-Dependent Activation of Late Replication Origins in S. cerevisiae. Molecular Cell 2, 173–182.

Donze, D., Adams, C.R., Rine, J., and Kamakaka, R.T. (1999). The boundaries of the silenced HMR domain in Saccharomyces cerevisiae. Genes Dev 13, 698–708.

Dorsett, D., and Merkenschlager, M. (2013). Cohesin at active genes: a unifying theme for cohesin and gene expression from model organisms to humans. Curr Opin Cell Biol 25, 327–333.

Feeney, K.M., Wasson, C.W., and Parish, J.L. (2010). Cohesin: a regulator of genome integrity and gene expression. Biochem J 428, 147–161.

Fernius, J., and Marston, A.L. (2009). Establishment of cohesion at the pericentromere by the Ctf19 kinetochore subcomplex and the replication fork-associated factor, Csm3. PLoS Genet 5, e1000629.

Fernius, J., Nerusheva, O.O., Galander, S., Alves Fde, L., Rappsilber, J., and Marston, A.L. (2013). Cohesin-dependent association of scc2/4 with the centromere initiates pericentromeric cohesion establishment. Curr Biol 23, 599–606.

Flick, J.S., and Johnston, M. (1990). Two systems of glucose repression of the GAL1 promoter in Saccharomyces cerevisiae. Mol Cell Biol 10, 4757–4769.

Fumasoni, M., and Murray, A.W. (2019). The evolutionary plasticity of chromosome metabolism allows adaptation to DNA replication stress. bioRxiv preprint.

Gagnon-Arsenault, I., Marois Blanchet, F.C., Rochette, S., Diss, G., Dube, A.K., and Landry, C.R. (2013). Transcriptional divergence plays a role in the rewiring of protein interaction networks after gene duplication. J Proteomics 81, 112–125.

Gay, S., Piccini, D., Bruhn, C., Ricciardi, S., Soffientini, P., Carotenuto, W., Biffo, S., and Foiani, M. (2018). A Mad2-Mediated Translational Regulatory Mechanism Promoting S-Phase Cyclin Synthesis Controls Origin Firing and Survival to Replication Stress. Mol Cell 70, 628–638 e625.

Gerstein, A.C., Chun, H.J., Grant, A., and Otto, S.P. (2006). Genomic convergence toward diploidy in Saccharomyces cerevisiae. PLoS Genet 2, e145.

Guacci, V., Koshland, D., and Strunnikov, A. (1997). A direct link between sister chromatid cohesion and chromosome condensation revealed through the analysis of MCD1 in S. cerevisiae. Cell 91, 47–57.

Heidinger-Pauli, J.M., Unal, E., Guacci, V., and Koshland, D. (2008). The kleisin subunit of cohesin dictates damage-induced cohesion. Mol Cell 31, 47–56.

Hinshaw, S.M., Makrantoni, V., Harrison, S.C., and Marston, A.L. (2017). The Kinetochore Receptor for the Cohesin Loading Complex. Cell 171, 72–84 e13.

Hinshaw, S.M., Makrantoni, V., Kerr, A., Marston, A.L., and Harrison, S.C. (2015). Structural evidence for Scc4-dependent localization of cohesin loading. Elife 4, e06057.

Hirano, T. (2012). Condensins: universal organizers of chromosomes with diverse functions. Genes Dev 26, 1659–1678.

Hittinger, C.T., and Carroll, S.B. (2007). Gene duplication and the adaptive evolution of a classic genetic switch. Nature 449, 677–681.

Ho, K.L., Ma, L., Cheung, S., Manhas, S., Fang, N., Wang, K., Young, B., Loewen, C., Mayor, T., and Measday, V. (2015). A role for the budding yeast separase, Esp1, in Ty1 element retrotransposition. PLoS Genet 11, e1005109.

Hottes, A.K., Freddolino, P.L., Khare, A., Donnell, Z.N., Liu, J.C., and Tavazoie, S. (2013). Bacterial adaptation through loss of function. PLoS Genet 9, e1003617.

Hu, B., Itoh, T., Mishra, A., Katoh, Y., Chan, K.L., Upcher, W., Godlee, C., Roig, M.B., Shirahige, K., and Nasmyth, K. (2011). ATP hydrolysis is required for relocating cohesin from sites occupied by its Scc2/4 loading complex. Curr Biol 21, 12–24.

Jerison, E.R., Kryazhimskiy, S., Mitchell, J.K., Bloom, J.S., Kruglyak, L., and Desai, M.M. (2017). Genetic variation in adaptability and pleiotropy in budding yeast. Elife 6.

Johnston, M., Flick, J.S., and Pexton, T. (1994). Multiple mechanisms provide rapid and stringent glucose repression of GAL gene expression in Saccharomyces cerevisiae. Mol Cell Biol 14, 3834–3841.

Klein, F., Mahr, P., Galova, M., Buonomo, S.B., Michaelis, C., Nairz, K., and Nasmyth, K. (1999). A central role for cohesins in sister chromatid cohesion, formation of axial elements, and recombination during yeast meiosis. Cell 98, 91–103.

Kohler, K., Sanchez-Pulido, L., Hofer, V., Marko, A., Ponting, C.P., Snijders, A.P., Feederle, R., Schepers, A., and Boos, D. (2019). The Cdk8/19-cyclin C transcription regulator functions in genome replication through metazoan Sld7. PLoS Biol 17, e2006767.

Koschwanez, J.H., Foster, K.R., and Murray, A.W. (2013). Improved use of a public good selects for the evolution of undifferentiated multicellularity. Elife 2, e00367.

Kryazhimskiy, S., Rice, D.P., Jerison, E.R., and Desai, M.M. (2014). Microbial evolution. Global epistasis makes adaptation predictable despite sequence-level stochasticity. Science 344, 1519–1522.

Kushnirov, V.V. (2000). Rapid and reliable protein extraction from yeast. Yeast 16, 857–860.

Laan, L., Koschwanez, J.H., and Murray, A.W. (2015). Evolutionary adaptation after crippling cell polarization follows reproducible trajectories. Elife 4.

Lang, G.I., Rice, D.P., Hickman, M.J., Sodergren, E., Weinstock, G.M., Botstein, D., and Desai, M.M. (2013). Pervasive genetic hitchhiking and clonal interference in forty evolving yeast populations. Nature 500, 571–574.

Lazar-Stefanita, L., Scolari, V.F., Mercy, G., Muller, H., Guerin, T.M., Thierry, A., Mozziconacci, J., and Koszul, R. (2017). Cohesins and condensins orchestrate the 4D dynamics of yeast chromosomes during the cell cycle. EMBO J 36, 2684–2697.

Lengronne, A., Katou, Y., Mori, S., Yokobayashi, S., Kelly, G.P., Itoh, T., Watanabe, Y., Shirahige, K., and Uhlmann, F. (2004). Cohesin relocation from sites of chromosomal loading to places of convergent transcription. Nature 430, 573–578.

Li, R., and Murray, A.W. (1991). Feedback control of mitosis in budding yeast. Cell 66, 519–531.

Li, Y., Muir, K.W., Bowler, M.W., Metz, J., Haering, C.H., and Panne, D. (2018). Structural basis for Scc3-dependent cohesin recruitment to chromatin. Elife 7.

Lundblad, V., and Zhou, H. (2001). Manipulation of plasmids from yeast cells. Curr Protoc Mol Biol Chapter 13, Unit13 19.

Makrantoni, V., Robertson, D., and Marston, A.L. (2019). Analysis of the Chromosomal Localization of Yeast SMC Complexes by Chromatin Immunoprecipitation. Methods Mol Biol 2004, 119–138.

Mangado, A., Morales, P., Gonzalez, R., and Tronchoni, J. (2018). Evolution of a Yeast With Industrial Background Under Winemaking Conditions Leads to Diploidization and Chromosomal Copy Number Variation. Front Microbiol 9, 1816.

Mantiero, D., Mackenzie, A., Donaldson, A., and Zegerman, P. (2011). Limiting replication initiation factors execute the temporal programme of origin firing in budding yeast. EMBO J 30, 4805–4814.

Marston, A.L. (2014). Chromosome segregation in budding yeast: sister chromatid cohesion and related mechanisms. Genetics 196, 31–63.

Mehta, G.D., Rizvi, S.M., and Ghosh, S.K. (2012). Cohesin: a guardian of genome integrity. Biochim Biophys Acta 1823, 1324–1342.

Melby, T.E., Ciampaglio, C.N., Briscoe, G., and Erickson, H.P. (1998). The symmetrical structure of structural maintenance of chromosomes (SMC) and MukB proteins: long, antiparallel coiled coils, folded at a flexible hinge. J Cell Biol 142, 1595–1604.

Michaelis, C., Ciosk, R., and Nasmyth, K. (1997). Cohesins: Chromosomal Proteins that Prevent Premature Separation of Sister Chromatids. Cell 91, 35–45.

Morgenthaler, A.B., Kinney, W.R., Ebmeier, C.C., Walsh, C.M., Snyder, D.J., Cooper, V.S., Old, W.M., and Copley, S.D. (2019). Mutations that improve the efficiency of a weak-link enzyme are rare compared to adaptive mutations elsewhere in the genome. bioRxiv preprint.

Nasmyth, K. (2002). Segregating sister genomes: the molecular biology of chromosome separation. Science 297, 559–565.

Nemet, J., Jelicic, B., Rubelj, I., and Sopta, M. (2014). The two faces of Cdk8, a positive/negative regulator of transcription. Biochimie 97, 22–27.

Nguyen Ba, A.N., Strome, B., Hua, J.J., Desmond, J., Gagnon-Arsenault, I., Weiss, E.L., Landry, C.R., and Moses, A.M. (2014). Detecting functional divergence after gene duplication through evolutionary changes in posttranslational regulatory sequences. PLoS Comput Biol 10, e1003977.

Orengo, C.A., and Thornton, J.M. (2005). Protein families and their evolution-a structural perspective. Annu Rev Biochem 74, 867–900.

Orgil, O., Matityahu, A., Eng, T., Guacci, V., Koshland, D., and Onn, I. (2015). A conserved domain in the scc3 subunit of cohesin mediates the interaction with both mcd1 and the cohesin loader complex. PLoS Genet 11, e1005036.

Paldi, F., Alver, B., Robertson, D., Schalbetter, S.A., Kerr, A., Kelly, D.A., Neale, M.J., Baxter, J., and Marston, A.L. (2019). Convergent genes shapes budding yeast pericentromeres. bioRxiv preprint.

Peric-Hupkes, D., and van Steensel, B. (2008). Linking cohesin to gene regulation. Cell 132, 925–928.

Rancati, G., Pavelka, N., Fleharty, B., Noll, A., Trimble, R., Walton, K., Perera, A., Staehling-Hampton, K., Seidel, C.W., and Li, R. (2008). Aneuploidy underlies rapid adaptive evolution of yeast cells deprived of a conserved cytokinesis motor. Cell 135, 879–893.

Roig, M.B., Löwe, J., Chan, K.-L., Beckouët, F., Metson, J., and Nasmyth, K. (2014). Structure and function of cohesin’s Scc3/SA regulatory subunit. FEBS Letters 588, 3692–3702.

Rosebrock, A.P. (2017). Synchronization and Arrest of the Budding Yeast Cell Cycle Using Chemical and Genetic Methods. Cold Spring Harb Protoc 2017.

Saayman, X., Ramos-Perez, C., and Brown, G.W. (2018). DNA Replication Profiling Using Deep Sequencing. Methods Mol Biol 1672, 195–207.

Schalbetter, S.A., Fudenberg, G., Baxter, J., Pollard, K.S., and Neale, M.J. (2019). Principles of meiotic chromosome assembly revealed in S. cerevisiae. Nat Commun 10, 4795.

Schindelin, J., Arganda-Carreras, I., Frise, E., Kaynig, V., Longair, M., Pietzsch, T., Preibisch, S., Rueden, C., Saalfeld, S., Schmid, B., et al. (2012). Fiji: an open-source platform for biological-image analysis. Nat Methods 9, 676–682.

Schleiffer, A., Kaitna, S., Maurer-Stroh, S., Glotzer, M., Nasmyth, K., and Eisenhaber, F. (2003). Kleisins: a superfamily of bacterial and eukaryotic SMC protein partners. Mol Cell 11, 571–575.

Schwob, E., and Nasmyth, K. (1993). CLB5 and CLB6, a new pair of B cyclins involved in DNA replication in Saccharomyces cerevisiae. Genes Dev 7, 1160–1175.

Shah, J.V., and Cleveland, D.W. (2000). Waiting for Anaphase. Cell 103, 997–1000.

Soppa, J. (2001). Prokaryotic structural maintenance of chromosomes (SMC) proteins: distribution, phylogeny, and comparison with MukBs and additional prokaryotic and eukaryotic coiled-coil proteins. Gene 278, 253–264.

Straight, A.F., Belmont, A.S., Robinett, C.C., and Murray, A.W. (1996). GFP tagging of budding yeast chromosomes reveals that protein–protein interactions can mediate sister chromatid cohesion. Current Biology 6, 1599–1608.

Straight, A.F., Marshall, W.F., Sedat, J.W., and Murray, A.W. (1997). Mitosis in living budding yeast: anaphase A but no metaphase plate. Science 277, 574–578.

Sunshine, A.B., Payen, C., Ong, G.T., Liachko, I., Tan, K.M., and Dunham, M.J. (2015). The fitness consequences of aneuploidy are driven by condition-dependent gene effects. PLoS Biol 13, e1002155.

Torres, E.M., Sokolsky, T., Tucker, C.M., Chan, L.Y., Boselli, M., Dunham, M.J., and Amon, A. (2007). Effects of aneuploidy on cellular physiology and cell division in haploid yeast. Science 317, 916–924.

Toth, A., Rabitsch, K.P., Galova, M., Schleiffer, A., Buonomo, S.B., and Nasmyth, K. (2000). Functional genomics identifies monopolin: a kinetochore protein required for segregation of homologs during meiosis i. Cell 103, 1155–1168.

Uhlmann, F., Lottspeich, F., and Nasmyth, K. (1999). Sister-chromatid separation at anaphase onset is promoted by cleavage of the cohesin subunit Scc1. Nature 400, 37–42.

Uhlmann, F., and Nasmyth, K. (1998). Cohesion between sister chromatids must be established during DNA replication. Current Biology 8, 1095–1102.

Uhlmann, F., Wernic, D., Poupart, M.A., Koonin, E.V., and Nasmyth, K. (2000). Cleavage of cohesin by the CD clan protease separin triggers anaphase in yeast. Cell 103, 375–386.

Voordeckers, K., Brown, C.A., Vanneste, K., van der Zande, E., Voet, A., Maere, S., and Verstrepen, K.J. (2012). Reconstruction of ancestral metabolic enzymes reveals molecular mechanisms underlying evolutionary innovation through gene duplication. PLoS Biol 10, e1001446.

Voordeckers, K., Kominek, J., Das, A., Espinosa-Cantu, A., De Maeyer, D., Arslan, A., Van Pee, M., van der Zande, E., Meert, W., Yang, Y., et al. (2015). Adaptation to High Ethanol Reveals Complex Evolutionary Pathways. PLoS Genet 11, e1005635.

Wildenberg, G.A., and Murray, A.W. (2014). Evolving a 24-hr oscillator in budding yeast. Elife 3.

Wu, N., and Yu, H. (2012). The Smc complexes in DNA damage response. Cell Biosci 2, 5.

Yona, A.H., Manor, Y.S., Herbst, R.H., Romano, G.H., Mitchell, A., Kupiec, M., Pilpel, Y., and Dahan, O. (2012). Chromosomal duplication is a transient evolutionary solution to stress. Proc Natl Acad Sci U S A 109, 21010–21015.

